# Defective Microhomology-Mediated End-joining in SMARCB1-Deficient Tumors

**DOI:** 10.1101/2025.11.20.689563

**Authors:** Guangli Zhu, Shuhei Asada, Huy Nguyen, Yuna Hirohashi, Lifang Sun, Avneesh Saravanapavan, Sirisha Mukkavalli, Geoffrey I. Shapiro, Alan D. D’Andrea

## Abstract

Rhabdoid tumors (RTs) are highly aggressive cancers driven by biallelic mutation of *SMARCB1,* a core subunit of the BAF (SWI/SNF) complex. We found that SMARCB1-deficient tumors have a defect in the microhomology-mediated end-joining (MMEJ) pathway, and SMARCB1 is essential for maintaining the protein level of the core MMEJ protein Polymerase Q (POLQ). Mechanistically, SMARCB1 facilitates the nuclear export of the *POLQ* mRNA through its interaction with the nuclear pore complex. Interestingly, loss of MMEJ in RT cells leads to a compensatory activation of, and a hyper-dependence on, the Fanconi Anemia (FA)/BRCA pathway. Knockout or inhibition of this pathway selectively kills RT cells. Notably, RBM39 degraders, novel splicing modulators, effectively inhibit the FA/BRCA pathway and kill RT cells. SMARCB1 and other cBAF/pBAF components are important for maintenance of MMEJ activity and POLQ protein level, suggesting that BAF-deficient cancers more broadly may be treated by targeted inhibition of the FA/BRCA pathway.

**Teaser:** SMARCB1-deficient tumors lose POLQ protein and MMEJ repair, forcing a hyper-dependence on the FA/BRCA pathway.

## INTRODUCTION

Rhabdoid tumors (RTs) are among the most aggressive and lethal types of pediatric cancers (*1*). These tumors are primarily driven by the biallelic inactivation of *SMARCB1*, an essential core subunit of the BAF (SWI/SNF) chromatin remodeling complex—specifically its canonical BAF (cBAF) and polybromo-associated BAF (pBAF) assemblies (*2*). Loss of SMARCB1 disrupts BAF complex integrity and function (*3, 4*), establishing it as the defining genetic alteration in RT. Despite intensive multimodal treatments including surgery, chemotherapy, and radiotherapy, the prognosis remains exceptionally poor, with 5-year survival rates often below 40% (*5*). Targeted therapies represent a promising treatment strategy of RTs, as illustrated by EZH2 inhibitors, which have received FDA approval for SMARCB1-negative malignancies (*6, 7*). Nonetheless, the therapeutic landscape for these tumors remains very limited, and the development of additional molecularly targeted strategies is urgently needed.

Unlike other high-grade cancers, RTs exhibit a remarkably simple genome, characterized by an extremely low mutation burden and minimal genomic instability, with biallelic *SMARCB1* loss representing the only recurrent genetic alteration (*8–11*). This genomic simplicity is paradoxical, given the aggressive nature of the disease and its sensitivity to specific agents targeting DNA damage repair proteins, such as ATR inhibition (*12*), although the mechanism of this drug sensitivity is unknown. The low mutation rate, coupled with the high sensitivity of RTs to replication stress, suggests a distinct defect in an error-prone DNA repair. We recently reported that, under replication stress, the loss of the microhomology-mediated end-joining (MMEJ) pathway sensitizes cells to ATR inhibitors (*13*). Therefore, RTs may have a defect in MMEJ, which would offer an opportunity for using inhibitors of other DNA repair pathways, via synthetic lethality.

DNA double-strand breaks (DSBs) are repaired by three distinct DNA repair pathways, including the nonhomologous end-joining (NHEJ) pathway, the HR pathway, and the MMEJ pathway (*14*). These pathways are known to function during the G1, S, or M phase of the cell cycle, respectively (*15–17*). The MMEJ pathway is driven primarily by the DNA Polymerase theta (POLQ) enzyme, a low-fidelity and error-prone polymerase which mediates microhomology-mediated repair of DNA DSBs, frequently resulting in characteristic mutations such as microdeletions or small insertions at DSB sites flanked by 2-20 bp homologous sequence (*18*). Other MMEJ proteins regulating the pathway include FEN1 (*19*), APEX2 (*19*), HMCES (*20*), XRCC1 (*21*), PARP1 (*22*) and LIG3 (*23*). The disruption of the MMEJ pathway is synthetic lethal with suppression of the NHEJ pathway and the HR pathway (*24–26*). Accordingly, NHEJ inhibitors (e.g., peposertib) and HR-pathway inhibitors (e.g., CDK12 inhibitors) are rational treatment strategies for MMEJ-deficient tumors (*24, 27*). More recently, POLQ/MMEJ-deficient cells were shown to be hypersensitive to perturbation of the Fanconi Anemia (FA)/BRCA pathway, which would be another promising treatment strategy for MMEJ-deficient tumors (*28, 29*).

The FA/BRCA pathway, comprising twenty-three FA genes, is a DNA damage repair pathway responsible for solving DNA interstrand crosslinks (*30*). FA/BRCA pathway-deficient ovarian tumors are hypersensitive to the genetic depletion or pharmacologic inhibition of the POLQ (*29*). Novobiocin, a known inhibitor of POLQ and the MMEJ pathway, shows strong *in vivo* efficacy in FA/BRCA pathway-deficient tumors. Similarly, a recent CRISPR screen revealed that disrupting the FA/BRCA pathway is synthetic lethal in cells with MMEJ defect (*28*). This evidence collectively supports FA/BRCA inhibition as a rational therapeutic approach for MMEJ-compromised tumors, motivating us to test whether SMARCB1-deficient RTs exhibit a similar dependency.

In the current report, we show that the CRISPR-mediated knockout of FA/BRCA genes resulted in a profound decrease in RT tumor cell viability. Since RT cells were dependent on the FA/BRCA pathway, we reasoned and confirmed that these cells are sensitive to inhibitors of the FA/BRCA pathway. Given that RT cells are driven by biallelic loss of the *SMARCB1* gene, we asked whether *SMARCB1* knockout would also increase cellular dependence on the FA/BRCA pathway. Indeed, knockout of *SMARCB1* in the RPE human cell line resulted in enhanced activation of the FA/BRCA pathway and an increased cellular dependence on FA/BRCA protein expression. Interestingly, *SMARCB1* loss resulted in a marked reduction in both MMEJ DNA repair activity and POLQ protein. Re-expression of *POLQ* cDNA restored MMEJ activity in SMARCB1-deficient cells, establishing POLQ insufficiency as the cause of the MMEJ defect in these cells. Mechanistically, SMARCB1 promotes POLQ protein expression primarily through its interaction with nuclear pore complex proteins that facilitate the export of the *POLQ* mRNA to the cytoplasm. Re-expression of wild-type (WT) SMARCB1, but not a SMARCB1 mutant defective in the interaction with other BAF subunits and the reassembly of the BAF complex, restored POLQ protein level and MMEJ activity. Taken together, we conclude that RT cells have a profound defect in the MMEJ pathway, caused by a lack of SMARCB1-mediated maintenance of POLQ protein level. The MMEJ defect in RT tumor cells accounts for their sensitivity to a broad range of DNA-damaging agents and for their vulnerability to inhibitors targeting the FA/BRCA pathway.

## RESULTS

### SMARCB1 deficiency drives dependence on and hyperactivation of the FA/BRCA pathway

Given the known DNA damage repair dysfunction in RTs, we anticipated a detectable shift in the genetic repair dependencies required for RT cell survival. To test this systematically, we initially interrogated the DepMap CRISPR screen dataset to identify genes whose inactivation by CRISPR/Cas9-mediated depletion selectively impaired the survival of RT cell lines (*31*), with a particular focus on the DNA damage repair genes. Interestingly, RT cell lines exhibited selective dependencies on multiple FA/BRCA genes (**Fig. 1A**). To validate this finding, individual FA/BRCA genes, including *FANCF*, *FANCA*, *FANCG*, and *BRCA2*, were knocked out by CRISPR/Cas9 in two RT cell lines (G401 and A204). Depletion of these genes, confirmed by immunoblot (**Fig. 1C**), markedly reduced clonogenic survival in RT cell lines (**Fig. 1B**). As expected, knockout of FANCA reduced FANCA and FANCG protein levels. Also, knockout of FANCG reduced FANCA and FANCG protein levels.

**Figure 1.**
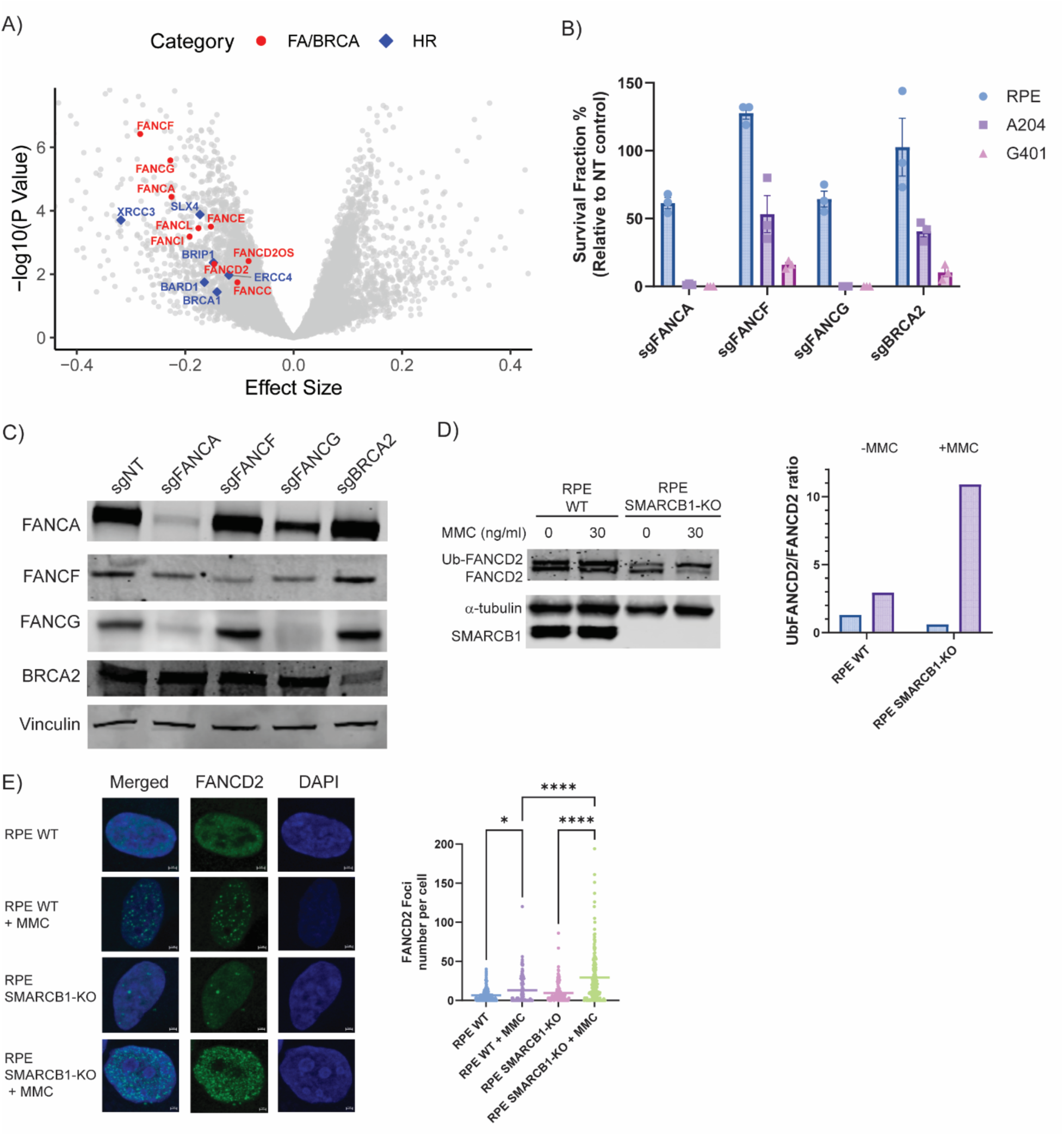
SMARCB1 deficiency drives dependence on and hyperactivation of the Fanconi Anemia/BRCA pathway. (A) Volcano plot showing DepMap CRISPR screen results highlighting significant hits in the Fanconi Anemia pathway. Genes in the FA/BRCA pathway are highlighted in red circles, while homologous recombination (HR) genes are highlighted in blue diamonds. (B) Clonogenic survival assays in RPE, G401, and A204 cells following expression of sgRNAs targeting non-targeting control (NT), *FANCF*, *FANCA*, *FANCG*, or *BRCA2*. (C) Immunoblot of FANCA, FANCF, FANCG, and BRCA2 in RPE *TP53^-/-^* cells expressing sgNT, sgFANCA, sgFANCF, sgFANCG, and sgBRCA2. Vinculin serves as a loading control. (D) Immunoblot of FANCD2, ubiquitinated FANCD2 (Ub-FANCD2), and SMARCB1 in RPE *TP53^-/-^* WT and *SMARCB1*-KO cells treated with or without MMC (30 ng/mL, 24 hours). α-tubulin serves as a loading control. Quantification of Ub-FANCD2/FANCD2 ratio is shown on the right. (E) Quantification (left) and representative immunofluorescence images (right) of FANCD2 nuclear foci formation in RPE *TP53^-/-^* WT and *SMARCB1*-KO cells with or without MMC treatment (30 ng/mL, 24 hours). Nuclei are stained with DAPI (scale bars = 2 μm). Data are shown as individual data points representing the number of FANCD2 foci per nucleus; the mean is indicated by a horizontal bar. Statistical significance was calculated using one-way ANOVA with Tukey’s post hoc test (*, *P* < 0.05; ****, *P* < 0.0001).

Since RTs are caused by biallelic loss-of-function mutations in the *SMARCB1* gene, we next determined whether the loss of the SMARCB1 protein causes the dysregulation of the FA/BRCA pathway in *SMARCB1*-deficient cells. We created an isogenic model by knocking out *SMARCB1* in human RPE cells using CRISPR/Cas9 and treated the cells with the DNA interstrand crosslinking agent mitomycin C (MMC). Interestingly, following treatment with MMC, we observed a significant activation of the FA/BRCA pathway in the *SMARCB1*-knockout RPE cells, as measured by monoubiquitination of FANCD2, a critical post-translational modification that signals the initiation of the repair cascade (**Fig. 1D**). The hyperactivation in the FA/BRCA pathway in *SMARCB1*-deficient cells was further confirmed by the elevated FANCD2 nuclear foci assembly following MMC treatment (**Fig. 1E**). Consistently, restoration of SMARCB1 in G401 cells reduces FANCD2 foci formation following MMC treatment (**Supplementary Fig. 1A-B**). These results indicate that the loss of SMARCB1 leads to a compensatory hyperactivation of the FA/BRCA pathway.

### SMARCB1 deficiency decreased MMEJ activity by reducing the POLQ protein level

We next evaluated the mechanism driving the FA/BRCA pathway dependency in *SMARCB1*-deficient cells. RT cell lines and *SMARCB1*-deficient RPE cells showed enhanced sensitivity to ATR inhibitors (**Supplementary Fig. 2A-B**), supporting the presence of intrinsic DDR defects. However, these cells are not sensitive to PARP inhibitors (**Supplementary Fig. 2C-D**), and data from the DepMap CRISPR screen further show that RT cell lines are dependent on the HR pathway for survival (**Fig. 1A**). Together, these findings indicate that RTs are not deficient in the HR pathway. Since recent studies have established a synthetic lethal relationship between the MMEJ pathway and the FA/BRCA pathway (*28, 29*), and RT cells also exhibit phenotypic features consistent with MMEJ defect, including decreased point mutagenesis and increased radiation sensitivity. Accordingly, we posited that SMARCB1 loss might inactivate MMEJ, thereby forcing RT cells to depend on the FA/BRCA signaling.

We therefore tested whether *SMARCB1* knockout would cause a defect in the MMEJ pathway (**Fig. 2A**). Using a CRISPR-based MMEJ reporter (*26*), we demonstrated that *SMARCB1* knockout results in a significant reduction in MMEJ activity. Since POLQ is the central component of the MMEJ pathway (*32, 33*), we initially tested whether *SMARCB1* knockout resulted in a loss of POLQ protein expression. Interestingly, knockout of *SMARCB1* resulted in a significant loss of the POLQ protein (**Fig. 2B**). However, knockout of *SMARCB1* did not affect the expression of other MMEJ proteins, including FEN1(*19*), LIG3(*23*), and HMCES(*20*), APEX2 (*19*), XRCC1 (*21*), and PARP1 (*22*) (**Supplementary Fig. 3**). Importantly, re-expression of WT SMARCB1 restored the POLQ protein level (**Fig. 2B**). To confirm that MMEJ defect was a direct consequence of POLQ depletion, we ectopically expressed WT *POLQ* in the SMARCB1-knockout cells, which can be successfully translated and was sufficient to fully rescue MMEJ activity (**Fig. 2C-D**). RT cell lines (G401 and A204) also exhibited reduced POLQ protein expression (**Fig. 2E**). Functionally, *SMARCB1* knockout also resulted in a loss of POLQ nuclear foci, indicating a failure to recruit POLQ to the sites of DNA double-strand breaks where its repair activity is required (**Fig. 2F**). Accordingly, the reduction in MMEJ activity following *SMARCB1* knockout is primarily due to the loss of POLQ. Furthermore, to assess POLQ/MMEJ activity in RT patients, we performed a mutational signature analysis in 56 primary RT patient samples from the NCI TARGET cohort using COSMIC v3.3 as the reference. The proportion of somatic SNVs attributed to single base substitution 3 (SBS3) signature, which is typically enriched in tumors with high POLQ (*9, 34*), is significantly lower in RTs (n = 56) compared with other pediatric malignancies, including Wilms Tumor (n = 81) and neuroblastoma (n = 135) (**Fig. 2G**).

**Figure 2.**
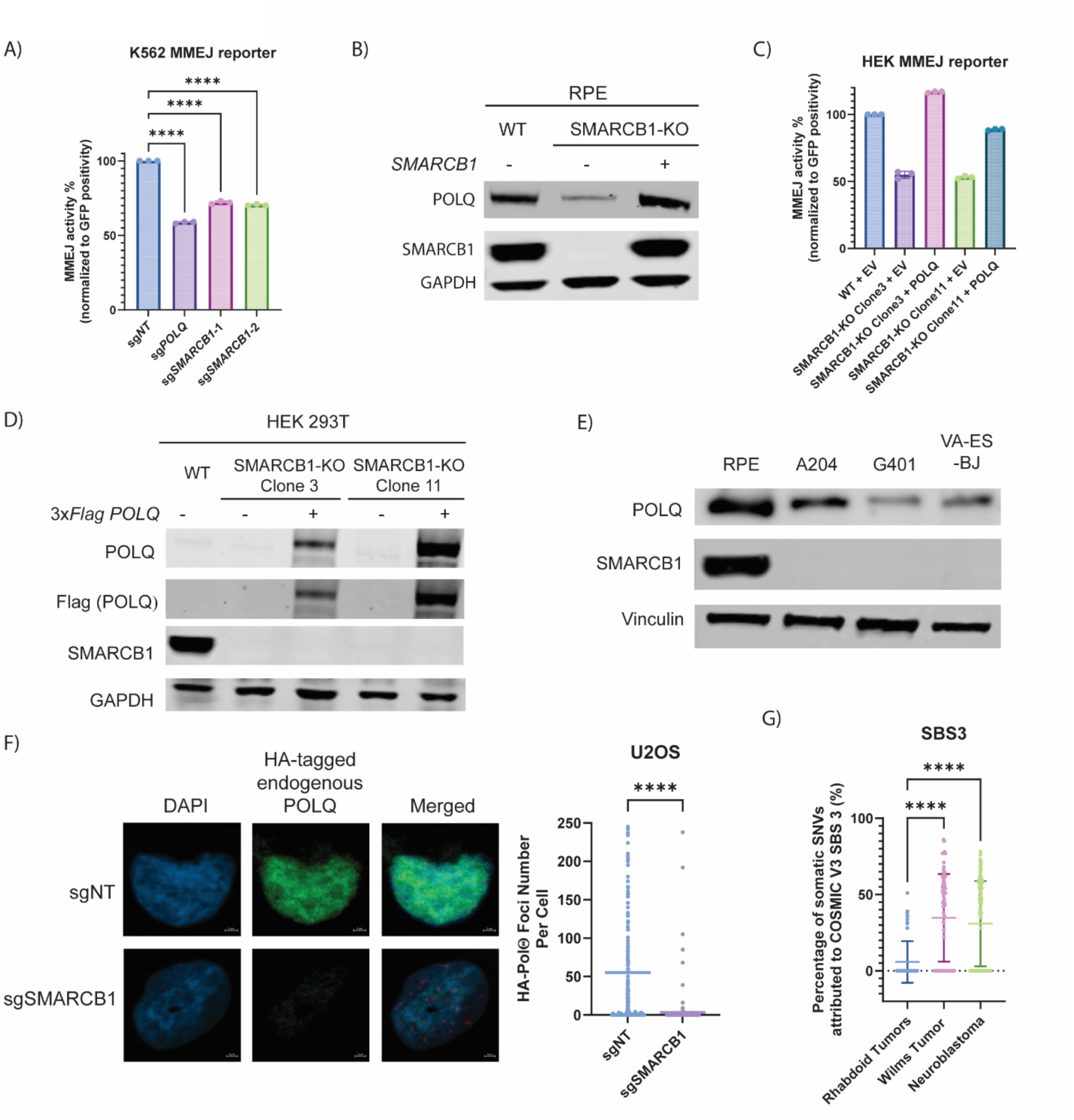
Loss of SMARCB1 confers POLQ-mediated MMEJ deficiency in rhabdoid tumors. (A) MMEJ reporter assay in K562 cells expressing the indicated sgRNAs. Data are presented as the mean ± SD. Statistical significance was calculated using one-way ANOVA with Tukey’s post hoc test (****, *P* < 0.0001). (B) Immunoblot of endogenous POLQ, SMARCB1, and GAPDH in WT RPE cells with empty vector, *SMARCB1*-KO RPE cells with empty vector, and *SMARCB1*-KO RPE cells reconstituted with WT SMARCB1. (C) MMEJ reporter assay in HEK293T WT and *SMARCB1*-KO cells with or without *3xFLAG POLQ* transfection. (D) Immunoblot of Flag-tagged POLQ expression in HEK293T WT and *SMARCB1*-KO cells with or without *3xFLAG POLQ* transfection. (E) Immunoblot of endogenous POLQ, SMARCB1, and GAPDH in WT RPE cells, *SMARCB1*-KO RPE cells, and rhabdoid tumor cell lines (G401, A204). (F) Representative immunofluorescence images showing nuclear foci of HA–tagged endogenous POLQ in U2OS cells expressing sg*NT* or sg*SMARCB1*. Quantification of HA– POLQ foci per cell is shown below. Data are shown as individual data points representing the number of HA-POLQ foci per nucleus; the mean is indicated by a horizontal bar. Statistical significance was calculated using Mann-Whitney U test (****, *P* < 0.0001). (G) Percentage somatic SNVs attributed to COSMIC single base substitution signatures (v3.3) for Rhabdoid Tumors (n = 56), Wilms Tumor (n = 81), and Neuroblastoma (n = 135). Data are shown as the mean ± SD. Statistical significance was calculated using one-way ANOVA with Tukey’s post hoc test (****, *P* < 0.0001).

Taken together, these data establish that the loss of SMARCB1 results in a profound dysfunction of the MMEJ pathway, which is primarily caused by the loss of the POLQ protein.

### SMARCB1 and other subunits of the cBAF/pBAF complex are required for maintaining POLQ expression and MMEJ function

SMARCB1 is a core subunit of the cBAF and pBAF chromatin remodeling complex (*4, 35*), and is important for the integrity of these complexes (**Supplementary Fig. 4A**). We therefore hypothesized that the broader BAF complex, and not just SMARCB1, is required for the maintenance of POLQ protein level and MMEJ activity. To investigate this, we individually knocked out selected subunits of the complex and tested their impact on MMEJ activity and POLQ protein level. As hypothesized, knockout of cBAF/pBAF subunits, including SMARCB1, SMARCA4, and a double knockout of ARID1A and ARID1B, resulted in a significant reduction in MMEJ activity (**Fig. 3A**). Correspondingly, knockout of these subunits also reduced the expression of POLQ, with SMARCB1 loss having the strongest effect **(Fig. 3B)**. To confirm that this function was specific to the cBAF/pBAF complexes, we knocked out BRD9, a subunit exclusive to the non-canonical BAF (ncBAF) complex. Strikingly, BRD9 loss had no impact on MMEJ activity or POLQ expression (**Fig. 3A, 3C, Supplementary Fig. 4B-C**). Moreover, pharmacological degradation of SMARCA4 using the degraders ACBI1 or BRM014 also led to reduced POLQ protein level and MMEJ activity (**Supplementary Fig. 4E-G**). These findings highlight a specific requirement for cBAF/pBAF subunits in maintaining POLQ expression and MMEJ function.

**Figure 3.**
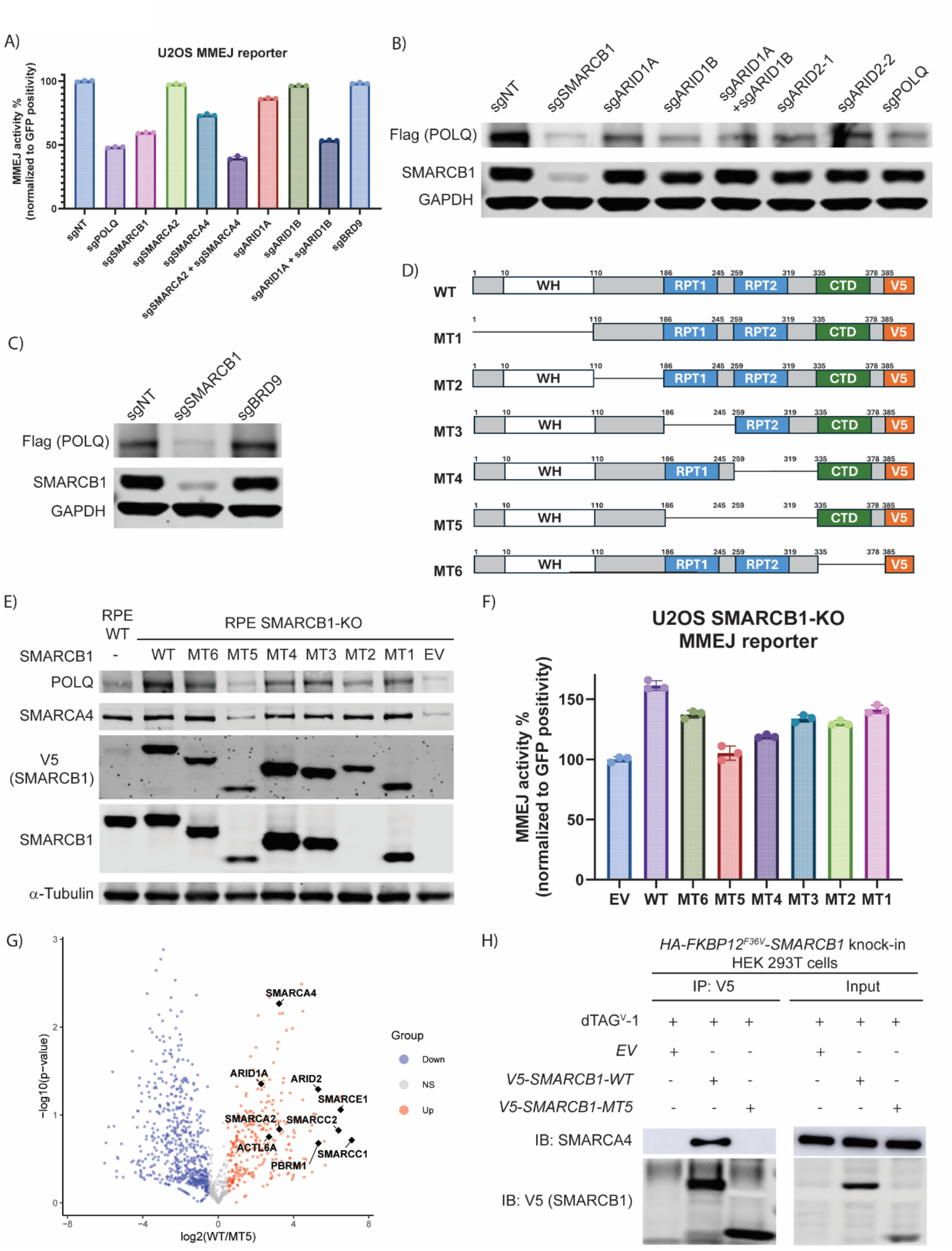
cBAF/pBAF integrity maintains POLQ levels and MMEJ activity. (A) MMEJ reporter assay in U2OS cells expressing the indicated sgRNAs. (B-C) Immunoblot analysis of Flag-tagged POLQ and SMARCB1 in RPE *TP53*^-/-^ *3xFLAG-POLQ* knock-in cells expressing indicated sgRNAs. GAPDH serves as a loading control. (D) Schematic of *SMARCB1* domain structure (WH: winged-helix, RPT1/2: repeat domains, CTD: C-terminal domain). SMARCB1 truncation mutants (MT1–MT6) are shown with their retained domains. (E) Immunoblot of indicated proteins in RPE *TP53*^-/-^ WT and *SMARCB1*-KO cells reconstituted with WT *SMARCB1* or truncation mutants. α-Tubulin serves as a loading control. (F) MMEJ reporter assay in U2OS *SMARCB1*-KO cells reconstituted with EV, WT SMARCB1, or truncation mutants (MT1–MT6). (G) Volcano plot showing proteins identified in WT-SMARCB1 versus MT5-SMARCB1 coimmunoprecipitates (n = 2). The x-axis represents the log₂ fold change (WT/MT5), and the y-axis represents the –log₁₀ p-value. Proteins enriched in WT (Up, red) or MT5 (Down, blue) are highlighted, whereas non-significant proteins are shown in grey. BAF complex subunits are marked as black-filled diamonds with white outlines and labeled with gene symbols. (H) Co-immunoprecipitation analysis. HEK293T *HA-FKBP12^F36V^-SMARCB1* knock-in cells were treated with dTAG-1 to induce endogenous SMARCB1 degradation and transfected with EV, V5-tagged WT SMARCB1, or V5-tagged MT5 mutant. Cell lysates were immunoprecipitated with anti-V5 antibody, followed by immunoblotting for SMARCA4 and V5 (SMARCB1).

Having established the role of the BAF complex, we next sought to determine the functional domain of SMARCB1 required for expression of the POLQ protein. We generated a series of SMARCB1 mutants with internal deletions of distinct regions (**Fig. 3D**). We next evaluated these mutant proteins for their ability to restore POLQ expression in a *SMARCB1*-knockout RPE cell line. Interestingly, the mutant 5 (MT5) of SMARCB1, which lacks both RPT1 and RPT2 domains, failed to restore POLQ expression (**Fig. 3E**). Consistent with this, among the SMARCB1 mutants, MT5 demonstrated the least ability to restore MMEJ activity in the MMEJ reporter assay (**Fig. 3F, Supplementary Fig. 4D**). We hypothesized that this phenotype arises from defective cBAF and pBAF assembly.

We next examined the binding partners of WT SMARCB1 and the MT5 SMARCB1 protein. Immunoprecipitation followed by mass spectrometry revealed that, unlike the WT SMARCB1, the MT5 mutant SMARCB1 protein failed to co-immunoprecipitate with other BAF subunits (**Fig. 3G, Supplementary Fig. 4H-I, Supplementary Table 1**). We validated this result by immunoprecipitation-western blotting and determined that the WT SMARCB1 binds to SMARCA4 while the mutant MT5 protein fails to co-immunoprecipitate with SMARCA4 (**Fig. 3H**). These findings are consistent with previous studies showing that RPT1 and RPT2 domains are required for the binding of SMARCB1 to other BAF subunits (*4*). Collectively, the RPT1 and RPT2 domains of SMARCB1 are essential for maintaining POLQ expression and MMEJ pathway function, and one essential function of these domains is to ensure BAF complex integrity, which is in turn required to maintain POLQ expression and MMEJ function.

### SMARCB1 maintains POLQ protein level primarily by promoting the nuclear export of *POLQ* mRNA to the cytoplasm

We next investigated the mechanism by which SMARCB1 expression regulates POLQ protein expression. Given BAF’s canonical role, we first assessed its impact on *POLQ* transcription. While SMARCB1 loss did lead to a statistically significant, yet modest, decrease in *POLQ* mRNA level (**Supplementary Fig. 5A**). However, the magnitude of this decrease was insufficient to account for the pronounced protein deficit. This critical disparity strongly indicated that while transcriptional control plays a supportive role, a more dominant, post-transcriptional mechanism must be the primary driver of POLQ regulation by SMARCB1. We next performed a series of validation experiments to systemically define the post-transcriptional mechanisms. First, cycloheximide chase assay revealed that POLQ protein stability was comparable between SMARCB1-KO cells and WT controls **(Supplementary Fig. 5B)**, and treatment with the proteasome inhibitor MG132 increases POLQ abundance to a similar degree in both groups, indicating no difference in proteasomal degradation **(Supplementary Fig. 5C)**. Second, actinomycin D chase experiment showed that *POLQ* mRNA stability was not affected by SMARCB1 status (**Supplementary Fig. 5D**). Finally, we performed deep RNA-seq for WT and SMARCB1-KO RPE *TP53*^-/-^ cells, and no substantial change in *POLQ* transcript was detected, except for a slight increase in intron 15 retention, which was confirmed by targeted RT-PCR (**Supplementary Fig. 5E-F)**. Finally, analysis from a previous study demonstrated that translation efficiency of *POLQ*, quantified as the ratio of ribosome-protected fragment density to total mRNA abundance, was not affected by SMARCB1 status (*36*), indicating that SMARCB1 loss does not alter the translational efficiency of *POLQ*.

Having systematically ruled out major effects on protein stability, mRNA stability, and translation, we re-interrogated our immunoprecipitation–mass spectrometry data, specifically comparing the interactomes of functional WT SMARCB1 versus the non-functional MT5 mutant. We found that WT SMARCB1 showed significantly stronger interaction with multiple Nuclear Pore Complex (NPC) proteins such as NUP153, NUP214, NUP188, and NUP133, compared to MT5 SMARCB1 mutant protein (**Fig. 4A, Supplementary Table 1**). In addition, nucleocytoplasmic transport ranked among the top enriched biological processes in the gene ontology analysis of proteins preferentially binding WT-SMARCB1 versus the MT5 mutant protein, further reinforcing this link (**Fig. 4B**). Consistent with these findings, co-immunoprecipitation revealed that WT SMARCB1 binds NUP214, NUP153, NUP188, NUP133, and NUP93 preferentially, while the MT5 mutant displayed markedly reduced interaction (**Fig. 4C**). Endogenous co-immunoprecipitations recapitulated this pattern, confirming the interaction at physiologic expression levels (**Fig. 4D**). A recent study demonstrated that the SMARCB1 protein may play a role in the regulation of mRNA export to the cytoplasm (*36*). In this study, long mRNAs were most strongly trapped in the nucleus by loss of SMARCB1.

**Figure 4.**
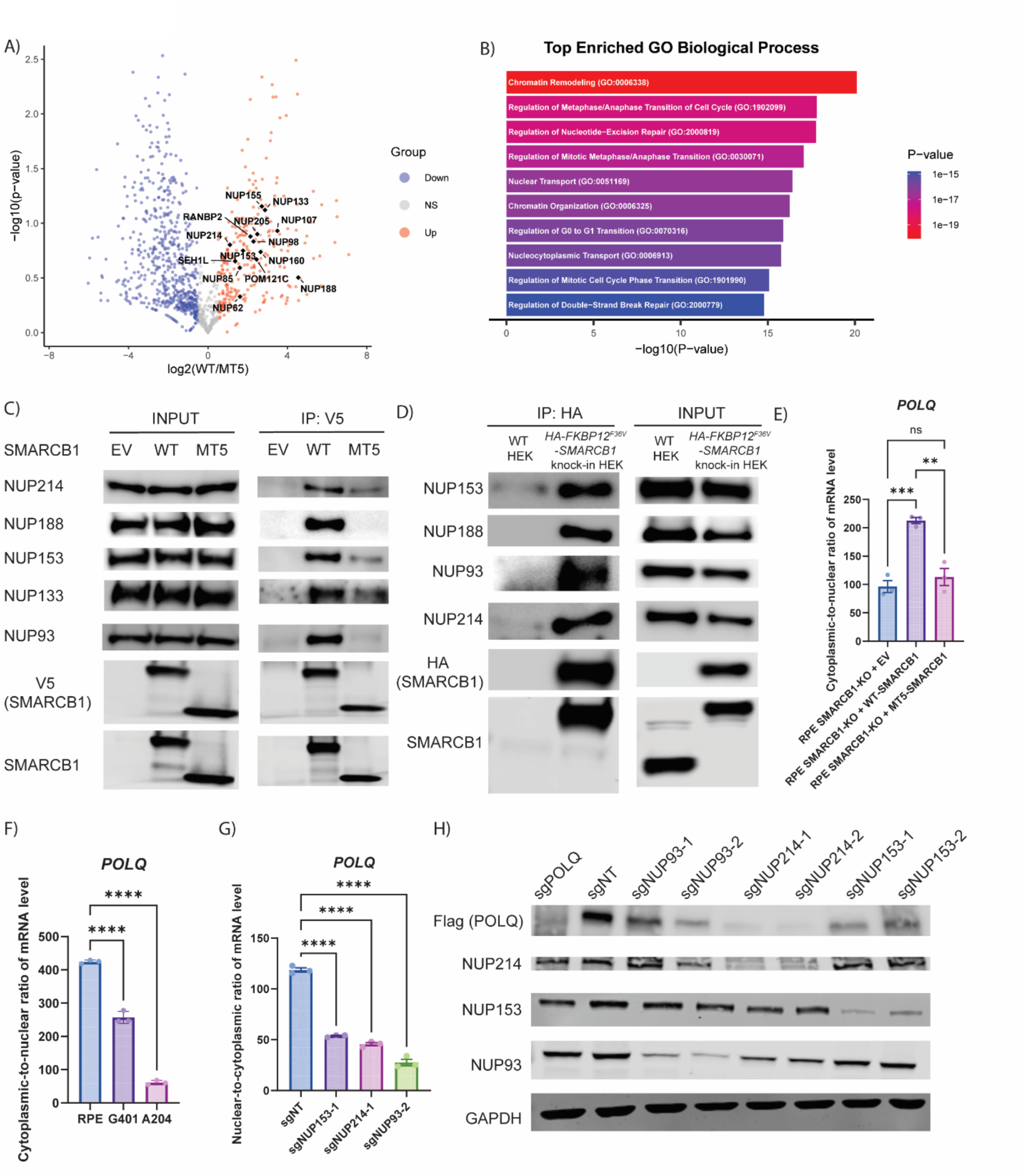
SMARCB1 maintains POLQ protein level by promoting nuclear export of its mRNA. (A) Volcano plot showing proteins identified in WT-SMARCB1 versus MT5-SMARCB1 coimmunoprecipitates (n = 2). The x-axis represents the log₂ fold change (WT/MT5), and the y-axis represents the –log₁₀ p-value. Proteins enriched in WT-SMARCB1 (Up, red) or MT5-SMARCB1 (Down, blue) are highlighted, whereas non-significant proteins are shown in grey. Nucleoporins are marked as black-filled diamonds with white outlines and labeled with gene symbols. (B) Gene Ontology enrichment analysis of proteins identified by mass spectrometry as enriched in WT-SMARCB1 versus MT5-SMARCB1 co-immunoprecipitates, showing the top biological processes. (C) Co-immunoprecipitation analysis. HEK293T *HA-FKBP12^F36V^-SMARCB1* knock-in cells were treated with dTAG-1 to induce endogenous SMARCB1 degradation and transfected with EV, V5-tagged WT-SMARCB1, or V5-tagged MT5-SMARCB1 mutant. Cell lysates were immunoprecipitated with anti-V5 antibody, followed by immunoblotting for NUP214, NUP188, NUP 153, NUP133, NUP93, SMARCB1, and V5. (D) Co-immunoprecipitation analysis. Cell lysates of WT HEK293T and *HA-FKBP12^F36V^-SMARCB1* knock-in HEK293T cells were immunoprecipitated with anti-HA antibody, followed by immunoblotting for NUP214, NUP188, NUP 153, NUP93, SMARCB1, and HA. (E) Quantification of cytoplasmic-to-nuclear mRNA ratios of *POLQ* in RPE *TP53*^−/−^ *SMARCB1*-KO cells reconstituted with EV, WT-SMARCB1, or MT5-SMARCB1, measured by qRT-PCR (n = 3). Data are shown as the mean ± SEM. Statistical significance was calculated using one-way ANOVA with Tukey’s post hoc test (****, *P* < 0.0001). (F) Quantification of cytoplasmic-to-nuclear mRNA ratios of *POLQ* and in *TP53*^−/−^ RPE, G401 and A204 cells, measured by qRT-PCR (n = 3). Data are shown as the mean ± SEM. Statistical significance was calculated using one-way ANOVA with Tukey’s post hoc test (****, *P* < 0.0001). (G) Quantification of cytoplasmic-to-nuclear mRNA ratios of *POLQ* and in *TP53*^−/−^ RPE expressing sgNT, sgNUP153-1, sgNUP214-1, sgNUP93-2, measured by qRT-PCR (n = 3). Data are shown as the mean ± SEM. Statistical significance was calculated using one-way ANOVA with Tukey’s post hoc test (****, *P* < 0.0001). (H) Immunoblot of POLQ, NUP153, NUP214, NUP93 and GAPDH in RPE *TP53*^-/-^ *3xFLAG-POLQ* knock-in cells expressing sgNT, sgPOLQ, sgNUP153, sgNUP214, and sgNUP93.

Since NPC proteins play a pivotal role in mRNA export (*37*), we hypothesized that SMARCB1 cooperates with NPC proteins to promote nuclear export of the *POLQ* mRNA, thereby ensuring its efficient translation and promoting MMEJ activity. As expected, *POLQ* mRNA export was severely impaired in both *SMARCB1*-deficient RPE cells and RT cell lines (**Fig. 4E-F**). Importantly, the ectopic expression of WT SMARCB1 but not the MT5 mutant SMARCB1 protein restored export of the *POLQ* mRNA in the *SMARCB1*-deficient RPE cells (**Fig. 4E**). Consistent with the mRNA export defect, knockout of NUP153, NUP214, or NUP93 led to a reduction of the cytoplasmic-to-nuclear ratio of *POLQ* mRNA and a marked decrease in POLQ protein level (**Fig. 4G-H**). Taken together, our data support a dual-function model for SMARCB1 in regulating POLQ. Beyond the canonical transcriptional role of the cBAF/pBAF complex, SMARCB1’s primary and indispensable function is to promote nuclear export of POLQ mRNA through its cooperation with the nuclear pore complex. The marked loss of POLQ protein in RTs thus reflects the combined failure of these SMARCB1-mediated processes.

### Rhabdoid tumors are vulnerable to FA/BRCA pathway inhibitors, including RBM39 degraders and CDK12 inhibitors

Taken together, these results demonstrate that SMARCB1 loss causes an MMEJ defect and a compensatory reliance on the FA/BRCA pathway, strongly suggesting that RT cells would be vulnerable to inhibitors of the FA/BRCA pathway. We first tested this hypothesis using recently identified small molecules that degrade the splicing factor RBM39, thereby inhibiting the FA/BRCA pathway by causing the loss of faithful splicing of mRNAs encoding key components in the FA/BRCA pathway (*26, 38*). As hypothesized, these RBM39 degraders, including E7820 and indisulam, induced selective lethality in the two RT cell lines (A204 and G401) and the SMARCB1-deficient epithelioid sarcoma cell line (VA-ES-BJ), while sparing the RPE control cell line (**Fig. 5A, 5D**). Consistent with these findings, both degraders selectively reduced the viability of U2OS SMARCB1-KO clones 4 and 7, whereas U2OS WT remained largely resistant across the tested concentrations (**Fig. 5B, 5E**). Importantly, the causal link between the BAF complex, MMEJ function, and this therapeutic dependency was also confirmed by the rescue experiments. Moreover, re-expression of WT SMARCB1 in U2OS SMARCB1-KO clone 4 attenuated sensitivity to both E7820 and indisulam, while an empty vector or the MT5 SMARCB1 mutant failed to rescue (**Fig. 5C, 5F**). A similar phenomenon was observed in the RT cell lines A204 and G401, further corroborating these findings (**Supplementary Fig. 6A-D**), underscoring that the structural integrity of the BAF complex is essential for maintaining MMEJ activity and preventing this synthetic lethal vulnerability. Immunoblot analysis confirmed that exposure of these cells to the RBM39 degraders caused a marked reduction in RBM39 protein and the FA/BRCA proteins, including FANCA and FANCD2, which in turn caused a significant increase in DNA double-strand breaks, as evidenced by elevated γ-H2AX levels (**Fig. 5G**).

**Figure 5.**
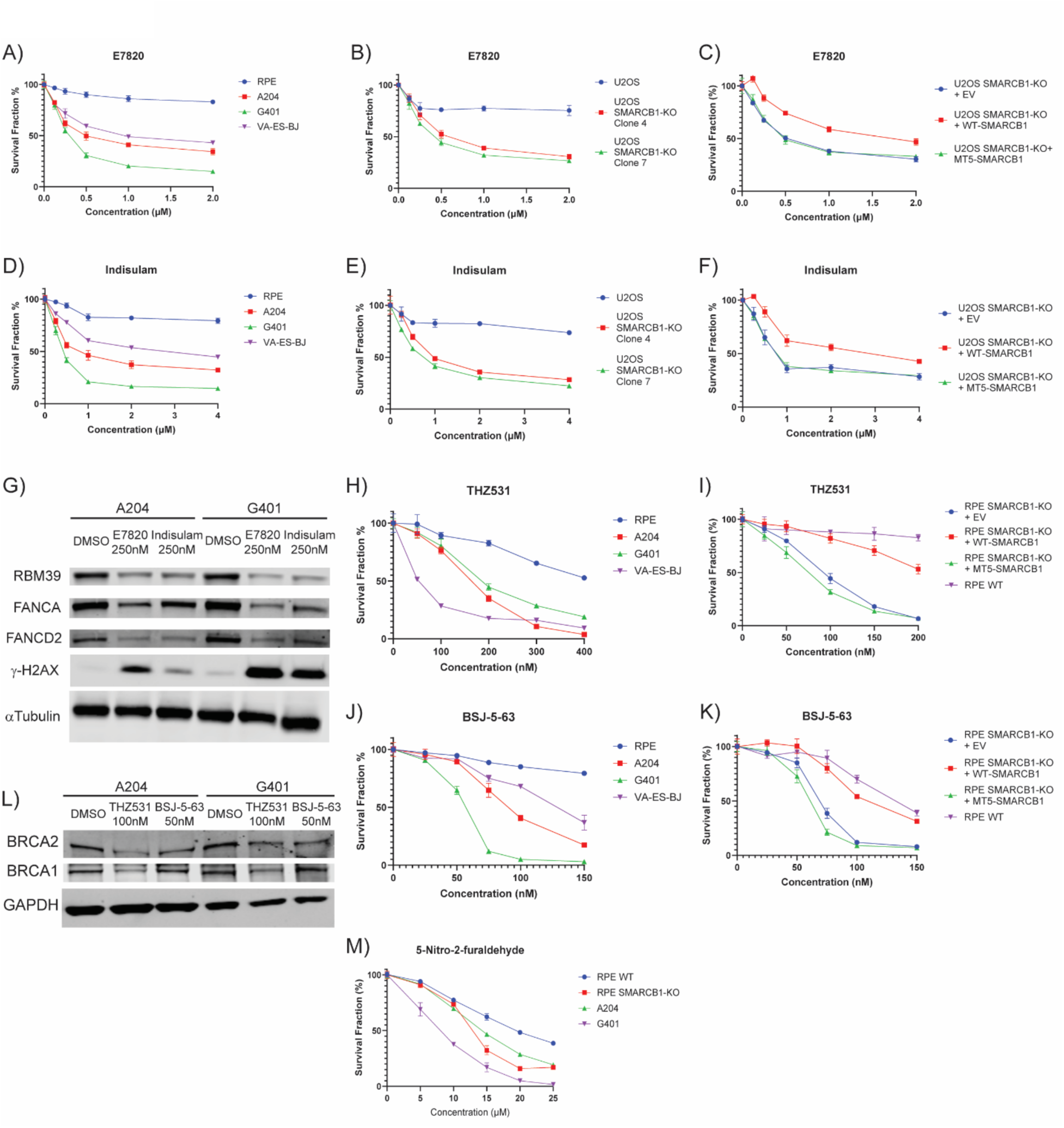
Rhabdoid tumors are vulnerable to Fanconi anemia/BRCA pathway inhibitors, including RBM39 degraders and CDK12 inhibitors. (A) Survival curves of RPE *TP53^-/-^* WT, rhabdoid tumor cell lines (A204, G401), and epithelioid sarcoma cell line (VA-ES-BJ) treated with indicated concentrations of E7820. (B) Survival curves of U2OS WT and U2OS *SMARCB1*-KO clone 4 and 7 cells treated with indicated concentrations of E7820. (C) Survival curves of U2OS *SMARCB1*-KO cells overexpressing empty vector, WT-SMARCB1, or MT5-SMARCB1 and treated with indicated concentrations of E7820. (D) Survival curves of RPE *TP53^-/-^* WT, rhabdoid tumor cell lines (A204, G401), and epithelioid sarcoma cell line (VA-ES-BJ) treated with indicated concentrations of Indisulam. (E) Survival curves of U2OS WT and U2OS *SMARCB1*-KO clone 4 and 7 cells treated with indicated concentrations of Indisulam. (F) Survival curves of U2OS *SMARCB1*-KO cells overexpressing empty vector, WT-SMARCB1, or MT5-SMARCB1 and treated with indicated concentrations of Indisulam. (G) Immunoblot of RBM39, FANCA, FANCD2, and γ-H2AX in A204 and G401 cells treated with DMSO, 250 nM E7820 or Indisulam. α-Tubulin serves as a loading control. (H-I) Survival curves of (H) RPE *TP53^-/-^* WT, rhabdoid tumor cell lines (A204, G401), and epithelioid sarcoma cell line (VA-ES-BJ) and (I) RPE *TP53^-/-^* WT and *SMARCB1*-KO cells reconstituted with EV, WT-SMARCB1, or MT5-SMARCB1 treated with indicated concentrations THZ531. (J-K) Survival curves of (J) RPE *TP53^-/-^* WT, rhabdoid tumor cell lines (A204, G401), and epithelioid sarcoma cell line (VA-ES-BJ) and (K) RPE *TP53^-/-^* WT and *SMARCB1*-KO cells reconstituted with EV, WT-SMARCB1, or MT5-SMARCB1 treated with indicated concentrations BSJ-5-63. (L) Immunoblot of BRCA2 and BRCA1 in A204 and G401 cells treated with DMSO, 100nM THZ531, or 50nM BSJ-5-63. GAPDH serves as a loading control. (M) Survival curves of RPE *TP53^-/-^* WT and *SMARCB1*-KO cells and rhabdoid tumor cell lines (A204, G401) treated with indicated concentrations of 5-Nitro-2-furaldehyde.

To confirm this vulnerability with mechanistically distinct agents, we next tested inhibitors of CDK12, such as THZ531 and BSJ-5-63, which were also reported to indirectly impair transcription of *BRCA* and *FANCF* genes and disrupt the FA/BRCA pathway and the HR pathway (*27, 39*). RT cells (A204 and G401), and VA-ES-BJ showed high sensitivity to THZ531 and BSJ-5-63, while the RPE *TP53*^-/-^ WT control remained resistant (**Fig. 5H, 5J**). In RPE *TP53*^-/-^ SMARCB1-KO cells, reconstitution with WT SMARCB1 markedly reduced sensitivity to both THZ531 and BSJ-5-63, whereas empty vector and the MT5 SMARCB1 mutant failed to restore resistance to these inhibitors (**Fig. 5I, 5K**). These observations were recapitulated in A204 and G401 cells, which further validate the finding **(Supplementary Fig. 6E–H)**. Immunoblot analysis confirmed that treatment with THZ531 or BSJ-5-63 led to a reduction in the protein levels of key HR pathway components, most notably BRCA2 (**Fig. 5L**). Similarly, SMARCB1-deficient RPE cells and RT cell lines also demonstrated hypersensitivity to 5-Nitro-2-furaldehyde (**Fig. 5M**), which is also known to inhibit the FA/BRCA pathway transcriptionally (*40*).

## DISCUSSION

Despite their rapid progression and poor prognosis, RTs characteristically harbor unusually simple genomes compared with other pediatric cancers. In this study, we identified a deficiency of an error-prone DNA repair pathway in RTs, MMEJ, that explains why RT cells harbor an extremely low mutational burden while simultaneously displaying hypersensitivity to replication stress. We demonstrated that SMARCB1 is indispensable for the maintenance of the MMEJ pathway, primarily through its non-canonical function of facilitating nuclear export of *POLQ* mRNA to maintain its protein expression. Loss of SMARCB1 reduces MMEJ activity and enforces a compensatory dependence on the FA/BRCA pathway, thereby creating a therapeutic vulnerability that can be exploited by targeting the FA/BRCA pathway.

Beyond our previous identification of Ewing sarcoma as an MMEJ-deficient tumor (*26*), RTs represent the second malignancy exhibiting this unique repair deficiency. We confirmed the MMEJ status of RT through a systematic approach. First, using a CRISPR-based MMEJ reporter assay, we provided the first direct functional evidence that SMARCB1 loss indeed diminished MMEJ activity. Next, we identified the deficiency of the core MMEJ enzyme, POLQ, as the direct cause of the MMEJ defect in RTs. Finally, we further validated this finding by demonstrating that a large cohort of primary RT patient samples was universally deficient in the SBS3 mutational signatures. Our results therefore demonstrate that RTs are MMEJ-deficient.

Our study reveals that the integrity of cBAF/pBAF complexes is essential for the maintenance of POLQ protein level, a function that involves both canonical and non-canonical BAF activities. The canonical role of the BAF complex in transcriptional control is observed in our study, where SMARCB1 loss results in a decrease in *POLQ* mRNA level. However, this minor transcriptional effect alone is insufficient to explain the profound decrease in POLQ protein. This discrepancy points to a dominant post-transcriptional defect. We performed a series of experiments to exclude contributions from different post-translational mechanisms, including mRNA stability, protein stability, and translation, to the reduction in POLQ protein level, and we finally identified and validated the regulation of mRNA nuclear export as a primary mechanism by detecting NPC proteins in the immunoprecipitation-mass spectrometry of SMARCB1. These observations align with a recent report that SMARCB1 loss alters the cytoplasmic-nuclear distribution of specific mRNAs, especially the long mRNAs, although the precise underlying mechanism remained undefined (*36*). Our study further expands the functions of the BAF complex beyond a transcriptional regulator. Supporting this, another study also found that SMARCA4 interacts with several NPC proteins, which is consistent with our findings that SMARCA4 KO also decreases MMEJ activity and POLQ protein level (*41*).

As the major channel between nucleus and cytoplasm for mRNA transport, NPC is a macromolecular complex composed of approximately 35 distinct proteins called nucleoporins (NUPs) (*37*). Structurally, the core consists of an inner ring and two outer-ring scaffolds (cytoplasmic and nucleoplasmic), composed of NUP188, NUP133, NUP98, NUP93, etc. Numerous FG-repeat nucleoporins, such as NUP153, NUP 214, and NUP98, line the transport channel and form the selective permeability barrier that is important for the selective transport. Our study showed that SMARCB1 not only interacts with the scaffold NUPs, but also interacts with NUPs essential for selective export, suggesting that chromatin remodeling complexes and nuclear export machineries are directly coupled through specific protein–protein interfaces to ensure the efficient translocation of specific mRNAs. This concept is highly consistent with the “gene gating” hypothesis originally proposed by Blobel, which postulates that actively transcribed loci can be recruited to the NPC to couple transcription with mRNA export in a spatially organized manner (*42*). NPC is not merely a passive channel for nucleocytoplasmic transport but also an active regulatory hub for gene expression. For example, in yeast, TREX-2 (Transcription-Export 2) complex coordinates interactions between transcriptional activators like the SAGA complexes and the nuclear basket proteins (Mlp1, the TPR homologue) to anchor active genes to the nuclear pore, thereby physically and functionally coupling transcription with mRNA export (*43–45*). Recent work in humans also shows that acute degradation of the TPR protein simultaneously led to defects in mRNA export and a reduction in nascent transcription for a specific subset of genes (*46*). Similar to our observations, a previous study reported that in colon cancer cell lines, ELYS recruits the *MYC* oncogene’s super-enhancer to the nuclear pore complex via β-catenin, thereby markedly enhancing the rate of mRNA export rather than significantly increasing transcription, ultimately leading to elevated MYC protein levels (*47*).

The evolving therapeutic landscape for SMARCB1-deficient malignancies has involved several distinct approaches. One prominent strategy targets the downstream epigenetic consequences of BAF complex dysregulation by the use of EZH2 inhibitors. This approach aims to counteract the functional imbalance in histone methylation caused by SMARCB1 loss (*6, 7*). Another set of approaches in development focuses more directly on the BAF machinery itself. These approaches include targeting DCAF5 (*48*) to prevent the degradation of partially assembled BAF complexes and restore their residual function, and targeting BRD9 (*49*), a key subunit of the non-canonical BAF complex that becomes essential for the survival of cancer cells lacking SMARCB1. While these approaches are promising, they directly target the complex and adaptable BAF machinery. Accordingly, this strategy may be prone to acquired resistance through complex reassembly of the BAF complex, modification of targets, or activation of alternative pathways (*50*).

In contrast, the therapeutic strategy demonstrated by our work offers a distinct and potentially more advantageous treatment paradigm. Our approach is built upon the established principle of synthetic lethality between the MMEJ pathway and the FA/BRCA or HR pathways (*28, 29*). We discovered that SMARCB1 loss renders RTs MMEJ-deficient, and this tumor-specific vulnerability directly rationalizes a therapeutic strategy of exploiting this defect by targeting the FA/BRCA and HR pathways. This strategy may enhance tumor specificity and mitigate resistance mechanisms linked to direct BAF complex inhibition. We validated the dependence of RTs on the FA/BRCA pathway by genetic perturbation, and then we employed pharmacological agents such as RBM39 degraders and CDK12 inhibitors to demonstrate the therapeutic application of this vulnerability. These agents successfully phenocopied the lethality of genetic knockout, confirming that this vulnerability is druggable, which highlights the potential of repurposing these existing agents for RTs and motivates the development of next-generation inhibitors designed to achieve greater selectivity in the FA/BRCA pathway.

Several limitations of our study should be noted. First, on a mechanistic level, while we have uncovered that the interaction between SMARCB1 and various NPC proteins is critical for *POLQ* mRNA export, the precise molecular architecture of this interaction remains to be elucidated. Deeper biochemical and structural studies are required to define the exact mode of interaction and to determine whether other cofactors, such as RNA-binding proteins, are involved in forming a dynamic export complex. Second, while our study focused on *POLQ* as the primary SMARCB1-regulated transcript, it does not systematically address the broader mRNA export landscape. Consequently, the potential impact of SMARCB1 loss on other transcripts remains unexplored.

In conclusion, by mechanistically linking SMARCB1 to the nuclear export of *POLQ* mRNA, our study explains the MMEJ deficiency in RTs and identifies the resulting dependency on the FA/BRCA pathway as a potent therapeutic vulnerability. The success of RBM39 degraders and CDK12 inhibitors in our models provides robust preclinical validation for targeting this pathway in RTs. This work not only expands the known functions of the BAF complex to include post-transcriptional regulation but also provides a clear and compelling foundation for a new therapeutic strategy against cancers driven by BAF complex aberrations.

## MATERIALS AND METHODS

### Cell Lines and Culture Conditions

This study utilized a panel of human cell lines, including HEK293T (ATCC CRL-3216), immortalized retinal pigment epithelial cells (RPE WT and RPE *TP53*^−/−^), K562 (ATCC CCL-243), U2OS (ATCC HTB-96), and rhabdoid tumor cell lines G401 (ATCC CRL-1441), A204 (ATCC HTB-82), and VA-ES-BJ (ATCC CRL-2138). All cell lines were cultured under standard conditions at 37°C in a humidified atmosphere with 5% CO₂. DMEM medium (Gibco, #11965092) was used for HEK293T, U2OS, and VA-ES-BJ cells, and DMEM F12 medium (Gibco, #10565042) was used for RPE cells. RPMI-1640 medium (Gibco, #11875119) was used for K562 cells, and McCoy’s medium (Gibco, #16600082) was used for G401 and A204 cells. All culture media were supplemented with 10% fetal bovine serum (Sigma Aldrich, #F2442) and a 1% penicillin-streptomycin solution (Gibco, #15140163).

### Compounds

The chemical compounds used in this study were as follows: MG132 (Selleck, #S2619), Cycloheximide (Selleck, #S7418), Actinomycin D (Selleck, #S8964), Mitomycin C (Selleck, #S8146), Olaparib (Selleck, #S1060), VE822 (Selleck, #S7102), ART558 (MedChem Express, #HY-141520), ACBI1 (Selleck, #S9612), BRM014 (Focus Biomolecules, #10-5244), Indisulam (Selleck, #S9742), E7820 (MedChem Express, #HY-14571), 5-Nitro-2-furaldehyde (Sigma-Aldrich, #170968), BSJ-5-63 (MedChem Express, #HY-162706), THZ531 (MedChem Express, #HY-103618), dTAGV-1 (Tocris, #6914), Polybrene (Sigma Aldrich, #TR-1003), Blasticidin (Life Technologies, #A1113903), Puromycin (Sigma Aldrich, #P7255). Stock solutions of all compounds were prepared in DMSO, stored in aliquots at -80°C, and diluted to final working concentrations immediately prior to use.

### Plasmid Constructs

psPAX2 and pMD2.G (Addgene #12260 and #12259) were gifts from Dr. Didier Trono. N106-EV-blast and N106-V5-BAF47-blast were gifts from Dr. Cigall Kadoch. For overexpression, complementary DNAs (cDNAs) encoding WT *SMARCB1*, various truncated mutants (MT1–MT6), and *POLQ* were subcloned into N106 or pcDNA3 expression vectors. The resultant proteins were engineered to contain FLAG, V5 or HA epitope tags to facilitate their detection and subsequent immunoprecipitation. To generate the plasmids mentioned above, site-directed mutagenesis was performed by standard techniques.

### Plasmid Transfection

Plasmid transfection was performed using Lipofectamine LTX with PLUS Reagent (Invitrogen, #15338100). Cells were seeded at 60–70% confluency one day prior to transfection. DNA–lipid complexes were prepared in Opti-MEM and added to the cells dropwise (Gibco #31985062). After 18 hours, the medium was replaced with fresh complete medium, and cells were harvested 48 hours – 72 hours post-transfection for downstream analysis.

### Viral Production and Transduction

Lentiviral particles were generated by transfecting HEK293T cells with a mixture of the lentiviral transfer plasmid, the packaging plasmid psPAX2, and the envelope plasmid pMD2.G using a calcium phosphate transfection kit (Takara Bio, #631312). The culture medium was replaced 18 hours post-transfection, and viral supernatants were harvested at 48hr, filtered through a 0.45 µm sterile filter, and used for infection. Target cells were transduced with virus in the presence of 8 µg/mL polybrene. Stably transduced cell populations were obtained by selection with puromycin or blasticidin.

### CRISPR/Cas9-Mediated Gene Editing

Gene knockouts were established using two methods. For stable knockouts, single-guide RNAs (sgRNAs) with sequences targeting specific genes were designed and subsequently cloned into the lentiCRISPRv2 vector via ligation using DNA Ligation Kit (Takara Bio, # 6023), and cells were transduced with lentiviruses co-expressing Cas9 and the specific sgRNA, followed by drug selection. Alternatively, ribonucleoprotein (RNP) complexes were formed by incubating recombinant Alt-R S.p. Cas9 Nuclease (Integrated DNA Technologies, #1081059) with synthetic recombinant sgRNA (Integrated DNA Technologies) and delivered into cells via electroporation using a 4D-Nucleofector system (Lonza). For endogenous epitope tag knock-in (e.g., 3x*FLAG*-*POLQ*, *HA*-*FKBP12*^F36V^-*SMARCB1*), RNP complexes were then delivered into cells transduced with virus expressing donor template via electroporation in the presence of Alt-R Cas9 Electroporation Enhancer (Integrated DNA Technologies, #1075915), and the media was changed the next day. Two days after electroporation, single-cell clones were subsequently isolated by limiting dilution in 96-well plates. Successful genomic editing in expanded clones was validated by immunoblot analysis and Sanger sequencing of the targeted locus. The sequences of all sgRNAs used in this study are provided in the Supplementary Methods.

### CellTiter-Glo Assay and Clonogenic Survival Assays

For evaluation of short-term cell viability, cells were seeded in 96-well plates and, on the following day, treated with a series of drug concentrations for 72–96 h. After treatment, luminescence—proportional to ATP content—was measured using the CellTiter-Glo Luminescent Cell Viability Assay (Promega, #G7573). For long-term survival, clonogenic assays were performed. Cells were seeded at low density in 6-well plates and exposed to the indicated drugs. After 7–14 days, cells were washed with PBS once, fixed with a mixture of 75% methanol and 25% acetate for 1 hour, stained with 0.5% crystal violet (Sigma-Aldrich, #548-62-9) in 20% methanol for 1 hour, and washed with water. After drying, the plates were imaged on GE Amersham Imager 600, and the area fraction of the colonies in each well were quantified using ImageJ software (National Institutes of Health, Bethesda, MD, USA, version 1.54). Both assays were performed in technical triplicates.

### Immunoblotting, Immunoprecipitation and Mass Spectrometry

For immunoblotting, whole-cell lysates were prepared by lysing cells in RIPA buffer (Cell Signaling Technology, #9806S) supplemented with protease and phosphatase inhibitor cocktails (Cell Signaling Technology, # 5872S) and PMSF (Cell Signaling Technology, #8553S) for 30 minutes on ice, centrifuging at 14000 rpm, 4℃ for 10 minutes, and collecting supernatants. Protein concentration was measured using the Bradford assay (Bio-Rad, #5000006). Protein samples were denatured by boiling at 70 °C for 10 minutes in 2× Laemmli sample buffer (Santa Cruz, #sc-286962). Equal protein amounts were separated on 3-8% NuPAGE Tris-Acetate gels (Invitrogen # EA0375BOX) or 4-12% Bis-Tris gels (Invitrogen # NP0321BOX) in NuPAGE Tris-Acetate SDS Running Buffer (Invitrogen, #LA0041) or NuPAGE MES SDS Running Buffer (Invitrogen, #NP0002), respectively, and transferred to nitrocellulose membranes (Bio-Rad, # 1620115). Membranes were blocked for 1 hour at room temperature with 5% milk (Bio-Rad, #1706404) in TBST (Tris-buffered saline with 0.1% Tween 20) and incubated with primary antibodies overnight at 4 °C. The membranes were incubated with either HRP-conjugated secondary antibodies (Cell Signaling Technology, #7076V, #7074S, #7077S) followed by incubating with ECL substrate (Life Technologies #34580) for chemiluminescent imaging on GE Amersham Imager 600, or DyLight 680/800-conjugated secondary antibodies (Cell Signaling Technology, #5470S or #5151S) for fluorescent detection on a LI-COR Odyssey CLx imaging system.

To prepare the nuclear extracts for co-immunoprecipitation, cells were harvested, washed with cold PBS, pelleted, and lysed in hypotonic buffer (50 mM Tris pH 7.5, 0.1% NP-40, 1 mM EDTA, 1 mM MgCl₂, protease inhibitors, 5μM MG132, and 1 mM PMSF) for 10 minutes with occasional flicking. After centrifugation at 14,000 rpm, 4 °C for 10 minutes, pelleted nuclei were resuspended in high-salt buffer (50 mM Tris pH 7.5, 300 mM NaCl, 1% NP-40, 1 mM EDTA, 1 mM MgCl₂, protease inhibitors, 5μM MG132, and 1 mM PMSF), and incubated on the agitator at 700rpm, 4 °C. After centrifugation at 14,000 rpm, 4 °C for 10 minutes, nuclear extracts were collected and supplemented with 1 mM DTT. For co-immunoprecipitation followed by mass spectrometry, HA-FKBP12^F36V^-SMARCB1 knock-in HEK 293T cells were transfected with the indicated plasmids (empty vector, pcDNA3-V5-SMARCB1 WT, or MT5), followed by treatment with 1 μM dTAGV-1 in fresh media the next day, and cells were harvested 24 hours after treatment.

For co-immunoprecipitation, 10% of the lysates were reserved as input, and the remaining lysates were incubated with primary antibodies pre-conjugated to Protein G magnetic beads (Invitrogen, #10004D) for 2 hours at 4 °C. The beads were then washed with washing buffer (20 mM Tris pH 7.5, 0.25% NP-40, 150 mM NaCl, protease inhibitors, 5μM MG132, and 1 mM PMSF) thrice and hypertonic buffer once. Proteins bound to the beads were eluted by boiling in 2x Laemmli sample buffer (Santa Cruz, #sc-286962) at 70 °C for 20 minutes and analyzed by immunoblotting or mass-spectrometry.

The detailed methods for mass-spectrometry was reported in our previous study (*26*). In brief, excised gel bands were subjected to in-gel digestion with sequencing-grade trypsin (Promega) overnight at 37°C. Peptides were extracted with 50% acetonitrile/1% formic acid, dried, and resuspended in 2.5% acetonitrile/0.1% formic acid. Samples were analyzed by nano-LC-MS/MS on an EASY-nLC system coupled to an Orbitrap Exploris480 mass spectrometer (Thermo Fisher Scientific). Peptides were separated on a C18 capillary column with a gradient of 90% acetonitrile/0.1% formic acid. Peptide sequences were identified using Sequest against a protein database, with a false discovery rate filtered to 1-2%. The resulting protein intensities were log2-transformed, normalized, and analyzed for differential abundance using the limma package in R. Volcano plots were generated using ggplot2 and ggrepel R packages.

### Immunofluorescence

Cells cultured on glass coverslips were subjected to treatments as specified. After treatment, cells were subsequently washed with PBS, fixed with 4% paraformaldehyde for 15 minutes at room temperature, and permeabilized with 0.3% Triton X-100 in PBS for 10 minutes at room temperature. Following blocking with 3% non-fat milk for 1 hour at room temperature, coverslips were incubated with primary antibodies diluted in blocking buffer overnight at 4 °C, washed with PBS thrice, and incubated with appropriate AlexaFluor488/568-conjugated secondary antibodies (Invitrogen, #A21202 or #A11031) diluted in blocking buffer for 1 hour at room temperature. After washing with PBS thrice, coverslips were mounted with ProLong Gold Antifade Mountant containing DAPI (Invitrogen, #P36934) and sealed with nail paint. Slides were scanned on Zeiss Axio Observer under 63X objective, and nuclear foci were quantified from at least 100 cells per condition using CellProfiler software (Broad Institute, Boston, MA, USA, version 4.2.6).

The information on primary antibodies used in this study is provided in the Supplementary Methods.

### MMEJ Reporter Assay

Establishment of CRISPR-based MMEJ reporter version 2 (V2) U2OS cells was reported in our previous study (*26*). 40,000 U2OS MMEJ reporter cells were seeded in 12-well plates, and the cells were transduced with lentivirus expressing Cas9, sgRNA targeting the reporter cassette, and BFP. For MMEJ reporter assay using HEK293T WT or SMARCB1-KO cells, these cells were transduced with the MMEJ reporter plasmid, the modified lentiCRISPRv2 plasmid expressing Cas9, sgRNA targeting the reporter cassette, and BFP, together with pcDNA3 EV or pcDNA3 POLQ WT cDNA. 72 hours after infection or transfection, the frequency of GFP-positive cells, which corresponds to the frequency of successful repair events, was determined by flow cytometry using CytoFLEX (Beckman). Flow cytometry data were analyzed using FlowJo (BD Biosciences, Milpitas, CA, version 10.8).

### RNA Analysis

Total RNA was extracted from cells using the RNeasy Mini Kit (Qiagen, #74104) and RNase-Free DNase Set (QIAGEN, #79254). RNA was reverse-transcribed into cDNA using the High-Capacity RNA-to-cDNA Kit (Applied Biosystems, #4368814). For quantitative analysis of transcript levels, qRT-PCR was performed using Power SYBR Green PCR Master Mix (Applied Biosystems, #4367659) and Quant Studio 7 flex Real-Time PCR System (Applied Biosystems), with expression values normalized to *GAPDH*.

Cytoplasmic and nuclear RNA were isolated using cytoplasmic and nuclear RNA purification kit (Norgen, #21000). For quantification of transcript abundance in each fraction, 30 ng of in vitro–transcribed *GFP* mRNA generated using HiScribe® T7 High Yield RNA Synthesis Kit (New England Biolabs, #E2040S) was added as a spike-in to 1 µg of cytoplasmic RNA and 1 µg of nuclear RNA, respectively, prior to reverse transcription. RNAs were co–reverse-transcribed to cDNA, and qRT-PCR was performed using Power SYBR™ Green PCR Master Mix (Thermo Fisher) with expression values normalized to the *GFP* spike-in.

To assess mRNA stability, cells were treated with the transcription inhibitor actinomycin D (5 µg/mL), and RNA levels were quantified at designated time intervals.

RT-qPCR assays were performed in technical triplicate. All PCR and RT-qPCR primers were listed in the supplementary methods.

### Mutational Signature Analysis

The somatic single nucleotide variants (SNVs) data of RTS, Neuroblastoma, and Wilms (WT) tumors generated by the Therapeutically Applicable Research to Generate Effective Treatments (TARGET) initiative were obtained from the Genomic Data Commons (https://portal.gdc.cancer.gov). The somatic SNVs were determined from the whole genome sequencing data of tumors and matched normal tissues. Profiles of SNV mutations were generated using SigProfilerMatrixGenerator (version 1.2.13; https://github.com/AlexandrovLab/SigProfilerMatrixGenerator). Mutational signature analysis was performed using SigProfilerAssignment (version 0.0.21; https://github.com/AlexandrovLab/SigProfilerAssignment). This involved fitting the mutational profiles to the COSMIC (https://cancer.sanger.ac.uk/signatures/) SNV signatures (SBS version 2).

### Statistical Analysis

Data was analyzed and visualized using GraphPad Prism (GraphPad Software, Boston, MA, version 10.4.1). Unless otherwise specified, all quantitative data are presented as the mean ± standard deviation (SD) or standard error of the mean (SEM) derived from a minimum of three independent experiments. The determination of statistical significance was performed using a two-tailed Student’s t-test or a Mann-Whitney U test for comparisons between two groups, or an analysis of variance (ANOVA) for multiple group comparisons. A p-value of less than 0.05 was established as the threshold for statistical significance.

## ACKNOWLEDGMENTS

We thank members of the D’Andrea laboratory for their helpful suggestions and comments. We thank Harvard Medical School and the Dana-Farber/Harvard Cancer Center in Boston, MA, for the use of the Taplin Mass Spectrometry core facilities.

## FUNDING

This work was supported by U.S. National Institutes of Health grants R01HL052725 and R01CA296618 (A.D.D.), the Breast Cancer Research Foundation (A.D.D.), the Ludwig Center at Harvard (A.D.D.), the David Liposarcoma Research Initiative at Harvard (A.D.D.), and the Smith Family Foundation (A.D.D.). This work was also supported by U.S. National Institutes of Health grants P50 CA168504 (A.D.D., G.I.S.), P50 CA240243 (A.D.D., G.I.S.), the SENSHIN Medical Research Foundation (S.A.), the Japan Society for the Promotion of Science Overseas Research Fellowships (S.A.).

## AUTHOR CONTRIBUTIONS

G.Z., S.A., and A.D.D. conceived the study. G.Z., S.A., Y.H., and L.S. designed and performed biological experiments and analyzed the data. H.N. and A.S. performed computational analysis. S.M. and G.I.S. provided scientific input. G.Z., S.A., S.M., and A.D.D. wrote the manuscript.

## COMPETING INTERESTS

A.D.D. reports consulting for AstraZeneca, Bayer AG, Bristol Myers Squibb, EMD Serono, GlaxoSmithKline, Impact Therapeutics, PrimeFour Therapeutics, Tango Therapeutics, Deerfield Management Company, Servier Bio-Innovation LLC, Roche Pharma, and Covant, Therapeutics; is an Advisory Board member for Impact Therapeutics; and reports receiving commercial research grants from Bristol Myers Squibb, EMD Serono, Moderna, and Tango Therapeutics.

## DATA AND MATERIALS AVAILABILITY

All data are available in the main text or the supplementary materials. The mass spectrometry data have been deposited to the ProteomeXchange Consortium via the PRIDE partner repository with the dataset identifier PXD069337. Raw RNA-Seq reads are available in the NCBI Sequence Read Archive under BioProject accession number PRJNA1344636. Source data are provided with this paper.

