## Supplementary Figures for "Defective Microhomology-Mediated End-joining in SMARCB1-Deficient Tumors"

#### Supplementary Text

sgRNA sequences

The sequences of single-guide RNAs (sgRNAs) used for CRISPR/Cas9-mediated gene knockout or targeting experiments are listed below.

| sgRNA | Sequences |
|---|---|
| *POLQ* | TCTGATCAATCGCCTCATAG |
| *SMARCB1-1* | GAGAACCTCGGAACATACGG |
| *SMARCB1-2* | ACAAGAGATACCCCTCACTC |
| *SMARCB1-8* | ATGATCCTACCTGAAGGCGT |
| *SMARCA4-1* | GCAGCGGGTACTCACGAGAG |
| *SMARCA4-2* | GCAGCAGACAGACGAGTACG |
| *SMARCC1* | TATTAGCTGATACCCCCTCT |
| *SMARCC2* | TATGCCTATTACCTGCACCA |
| *SMARCD1* | TATTAAGACACATAAGCTCC |
| *SMARCE1* | TATGTAAGCAAGGTACGCGG |
| *ARID1A* | AGCGGGATCAGGATCTATGC |
| *ARID1B* | CCCATGATGCGGAGCTACGG |
| *ARID2-1* | GAAGAAGATGGAGTTATGTT |
| *ARID2-2* | TGTGGTAGGAGTAAAACGGA |
| *BRD7-1* | GAAGTCACCGAACTCTCCAC |
| *BRD7-2* | TCGGACAAACACCTCTACGA |
| *BICRA* | ACCATCTGGAAGGTCCGAGG |
| *BRD9* | ATAATGCAATGACATACAAT |
| *FANCG* | GCCTACCTTGCAGACTATG |
| *FANCA* | GGGTATTCTCTCAGCCGGGA |
| *BRCA2* | CGCTTCCGCGGCCCGTTCAA |
| *NUP153-1* | TAATGCAGGGGATTCCAACA |
| *NUP153-2* | AAGGCAGACAAGTTGAATGC |
| *NUP93-1* | AGTACCTGTAGTCCAGGCAA |
| *NUP93-2* | AGGCTCAAGAGGCTCAAAGG |
| *NUP214-1* | ATACACTATGGCGAAGACGT |
| *NUP214-2* | ATAGTCTACGACAACTCCCA |

Antibodies Used for Immunoprecipitation, Immunofluorescence, and Immunoblot

A list of antibodies used for immunoprecipitation (IP), immunofluorescence (IF), and immunoblot, along with their sources, dilution ratio, clone information and catalog numbers, is provided below.

| IP Antibodies | Source | Clone | Identifier |
|---|---|---|---|
| HA | Roche | 3F10 | 11867423001 |
| V5 | Abcam | SV5-Pk1 | ab27671 |

| IF Antibodies | Source | Dilution | Clone | Identifier |
|---|---|---|---|---|
| HA | Cell Signaling Technology | 1:200 | C29F4 | 3724S |
| FANCD2 | Novus | 1:500 | Polyclonal | NB100182 |

| Immunoblot Antibodies | Source | Dilution | Clone | Identifier |
|---|---|---|---|---|
| **HA** | Cell Signaling Technology | 1:1000 | C29F4 | 3724S |
| **V5** | Cell Signaling Technology | 1:1000 | D3H8Q | 13202S |
| **V5** | Abcam | 1:1000 | SV5-Pk1 | ab27671 |

| | | | | |
|---|---|---|---|---|
| **Flag** | Sigma Aldrich | 1:1000 | M2 | F1804 |
| **GAPDH** | Cell Signaling Technology | 1:1000 | D16H11 | 5174S |
| **GAPDH** | Cell Signaling Technology | 1:1000 | D4C6R | 97166S |
| **α -Tubulin** | Cell Signaling Technology | 1:1000 | 11H10 | 2125S |
| **Vinculin** | Cell Signaling Technology | 1:1000 | Polyclonal | 4650S |
| **SMARCB1** | Cell Signaling Technology | 1:1000 | D8M1X | 91735S |
| **SMARCA4** | Cell Signaling Technology | 1:1000 | A52 | 3508S |
| **NUP214** | Abcam | 1:1000 | Polyclonal | ab70497 |
| **NUP153** | Cell Signaling Technology | 1:1000 | E3I6Z | 98559S |
| **NUP133** | Santa Cruz Biotechnology | 1:200 | E-6 | SC-376763 |
| **NUP93** | Santa Cruz Biotechnology | 1:1000 | E-8 | SC-374399 |
| **NUP188** | Bethyl Laboratories | 1:1000 | Polyclonal | A302322AT |
| **FANCD2** | Santa Cruz Biotechnology | 1:200 | FI17 | SC-20022 |
| **FANCA** | Cell Signaling Technology | 1:1000 | D1L2Z | 14657S |
| **FANCF** | Santa Cruz Biotechnology | 1:200 | D-2 | SC-271952 |
| **FANCG** | This study | 1:1000 | Polyclonal | |
| **BRCA2** | Sigma Aldrich | 1:1000 | 2B | OP95 |
| **BRCA1** | Cell Signaling Technology | 1:1000 | Polyclonal | 9010S |
| **RBM39** | Proteintech | 1:1000 | Polyclonal | 21339-1-AP |
| **γH2AX** | Cell Signaling Technology | 1:1000 | 20E3 | 9718S |
| **PARP1** | Abcam | 1:1000 | E102 | ab32138 |
| **LIG3** | Abcam | 1:1000 | Polyclonal | ab185815 |
| **XRCC1** | Cell Signaling Technology | 1:1000 | E4A3V | 76998S |
| **HMCES** | Atlas Antibodies | 1:1000 | Polyclonal | HPA044968 |
| **FEN1** | Cell Signaling Technology | 1:1000 | E4S8C | 82354S |
| **APEX2** | Cell Signaling Technology | 1:1000 | E5A2Z | 74728S |
| **POLQ** | Cell Signaling Technology | 1:1000 | 64708 | 64708 |

#### Primer Sequences

The nucleotide sequences of primers used for PCR amplification and quantitative real-time PCR (qRT-PCR) analyses are listed below.

| **PCR Primers** | **Forward Primer Sequence** | **Reverse Primer Sequence** |
|---|---|---|
| ***POLQ exon15*** | CTTTCCCAATTTCAGAAGCG | TCTGCCACAGTATGAAAGCC |
| ***GAPDH*** | GGAGCGAGATCCCTCCAAAAT | GGCTGTTGTCATACTTCTCATG G |
| ***POLQ Intron15-Exon16 Junction*** | CAGTTCCTGGTAACAGCTCAGA | TTGCGACGTTCTTCAACTGC |

| **qPCR Primers** | **Forward Primer Sequence** | **Reverse Primer Sequence** |
|---|---|---|
| ***POLQ*** | ACCTCTCCATCAAGGCATTTCT | GCAAAAGTTCCAGCAGATACCC |
| ***GAPDH*** | GGAGCGAGATCCCTCCAAAAT | GGCTGTTGTCATACTTCTCATGG |
| ***GFP*** | CGCGCTTCTCGTTGGGGTCT | AGCAGAACACCCCCATCGGC |

Next generation sequencing

RNA samples were prepared using RNeasy Mini Kit (Qiagen, #74104) and RNase-Free DNase Set (QIAGEN, #79254). Deep RNA-seq (300 million reads/sample, N = 3 for each cell line) was performed for RPE *TP53*$^{-/-}$ WT and SMARCB1-KO cells. RNA samples were quantified using Qubit 2.0 Fluorometer (ThermoFisher Scientific, Waltham, MA, USA) and RNA integrity was checked with 4200 TapeStation (Agilent Technologies, Palo Alto, CA, USA). Strand-specific RNA sequencing library was prepared by using NEBNext Ultra II Directional RNA Library Prep Kit for Illumina following manufacturer's instructions (NEB, Ipswich, MA, USA). Briefly, the enriched RNAs were fragmented for 8 minutes at 94 °C. First strand and second strand cDNA were subsequently synthesized. The second strand of cDNA was marked by incorporating dUTP during the synthesis. cDNA fragments were adenylated at 3'ends, and indexed adapter was ligated to cDNA fragments. Limited cycle PCR was used for library enrichment. The incorporated dUTP in second strand cDNA quenched the amplification of second strand, which helped to preserve the strand specificity. The sequencing library was validated on the Agilent TapeStation (Agilent Technologies, Palo Alto, CA, USA), and quantified by using Qubit 2.0 Fluorometer (ThermoFisher Scientific, Waltham, MA, USA) as well as by quantitative PCR (KAPA Biosystems, Wilmington, MA, USA). The sequencing libraries were multiplexed and clustered onto a flowcell on the Illumina NovaSeq instrument according to manufacturer's instructions. The samples were sequenced using a 2x150bp Paired End (PE) configuration. Image analysis and base calling were conducted by the NovaSeq Control Software (NCS). Raw sequence data (.bcl files) generated from Illumina NovaSeq was converted into fastq files and de-multiplexed using Illumina bcl2fastq 2.20 software. One mis-match was allowed for index sequence identification.

### Supplementary Figure 1

A)

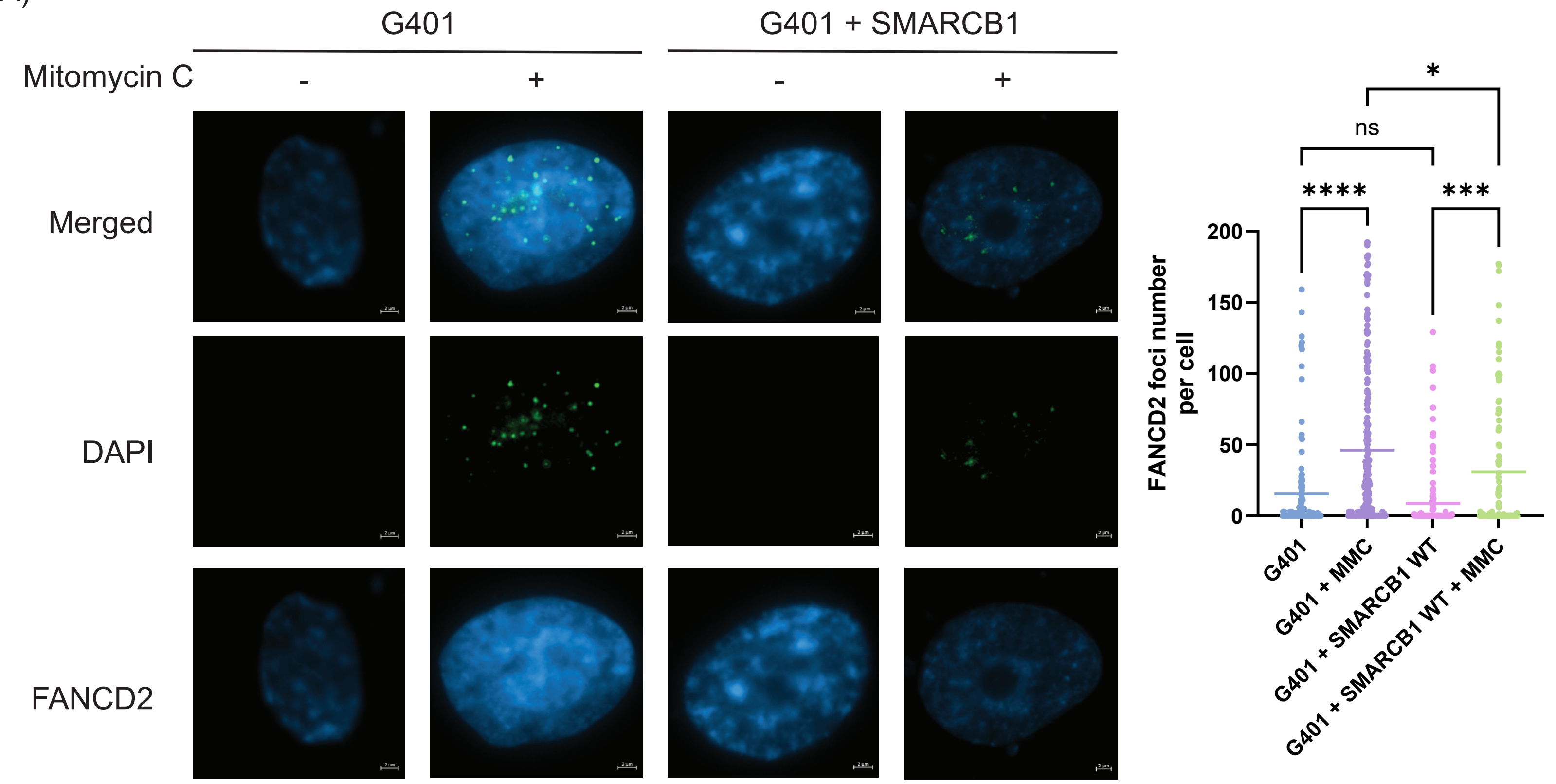


B)

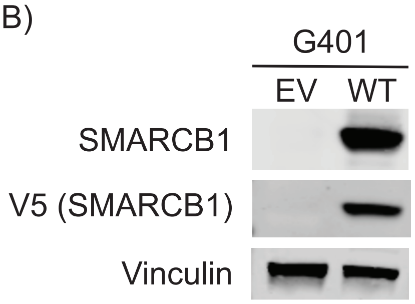

**Fig. S1. SMARCB1 restoration reduces FANCD2 foci formation in G401 cells.**

(A) Representative immunofluorescence images (left) and quantification (right) of FANCD2 nuclear foci formation in wild-type and SMARCB1-complemented G401 cells with or without MMC treatment. Nuclei are stained with DAPI (scale bars = 2μm). Data are shown as individual data points representing the number of FANCD2 foci per nucleus; the mean is indicated by a horizontal bar. Statistical significance was calculated using one-way ANOVA with Tukey's post hoc test (*, $P < 0.05$; ***, $P < 0.001$; ****, $P < 0.0001$).

(B) Immunoblot of SMARCB1, V5, and Vinculin in G401 cells expressing empty vector and V5-tagged SMARCB1.

### Supplementary Figure 2

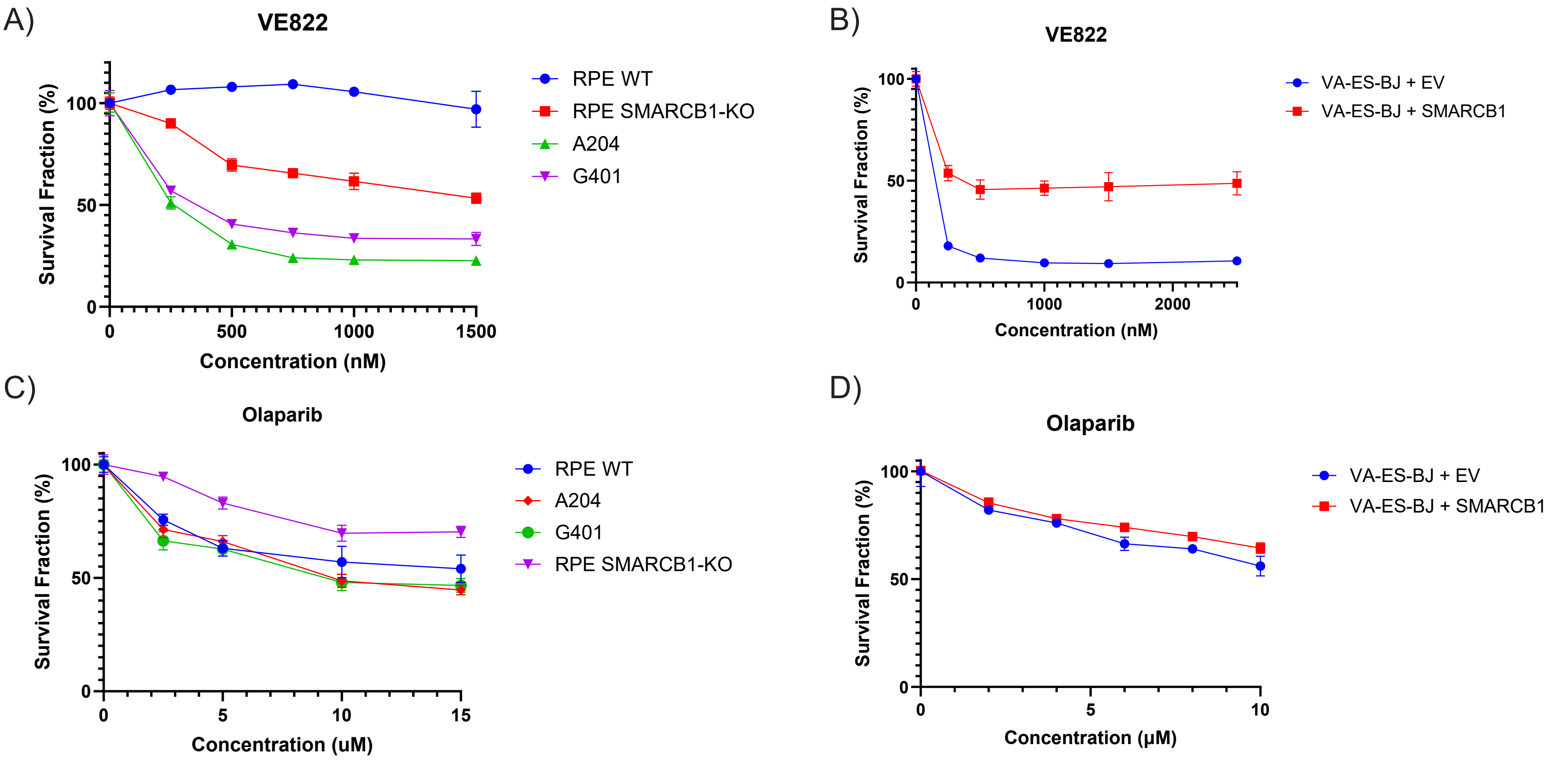

**Fig. S2. SMARCB1-deficient cells and rhabdoid tumor cell lines are sensitive to replication stress, but not PARP inhibition.**

(A) Survival curves of RPE *TP53*$^{-/-}$ WT, *SMARCB1*-KO, A204, and G401 cells treated with the ATR inhibitor VE822.

(B) Survival curves of VA-ES-BJ cells reconstituted with EV or SMARCB1 and treated with VE822.

(C) Survival curves of RPE *TP53*$^{-/-}$ WT, *SMARCB1*-KO, and rhabdoid tumor cell lines (A204, G401) treated with indicated concentrations of Olaparib.

(D) Survival curves of VA-ES-BJ cells transduced with EV or SMARCB1 and treated with Olaparib.

### Supplementary Figure 3

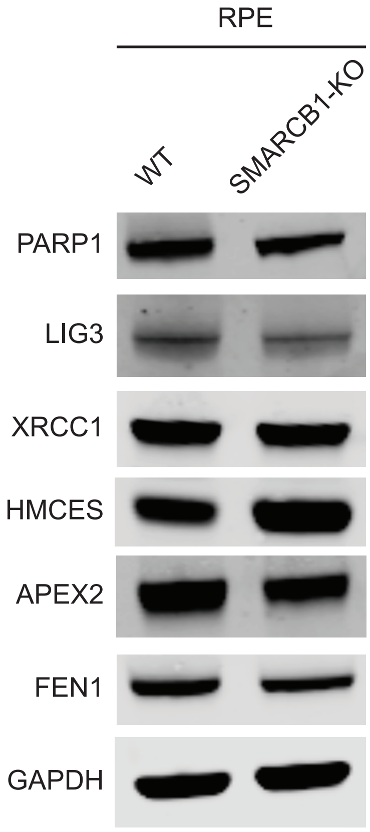

**Fig. S3. Expression of MMEJ pathway proteins in *SMARCB1*-deficient cells.**

(A) Immunoblot of indicated proteins involved in MMEJ pathways in RPE *TP53*$^{-/-}$ WT and *SMARCB1*-KO cells.

### Supplementary Figure 4

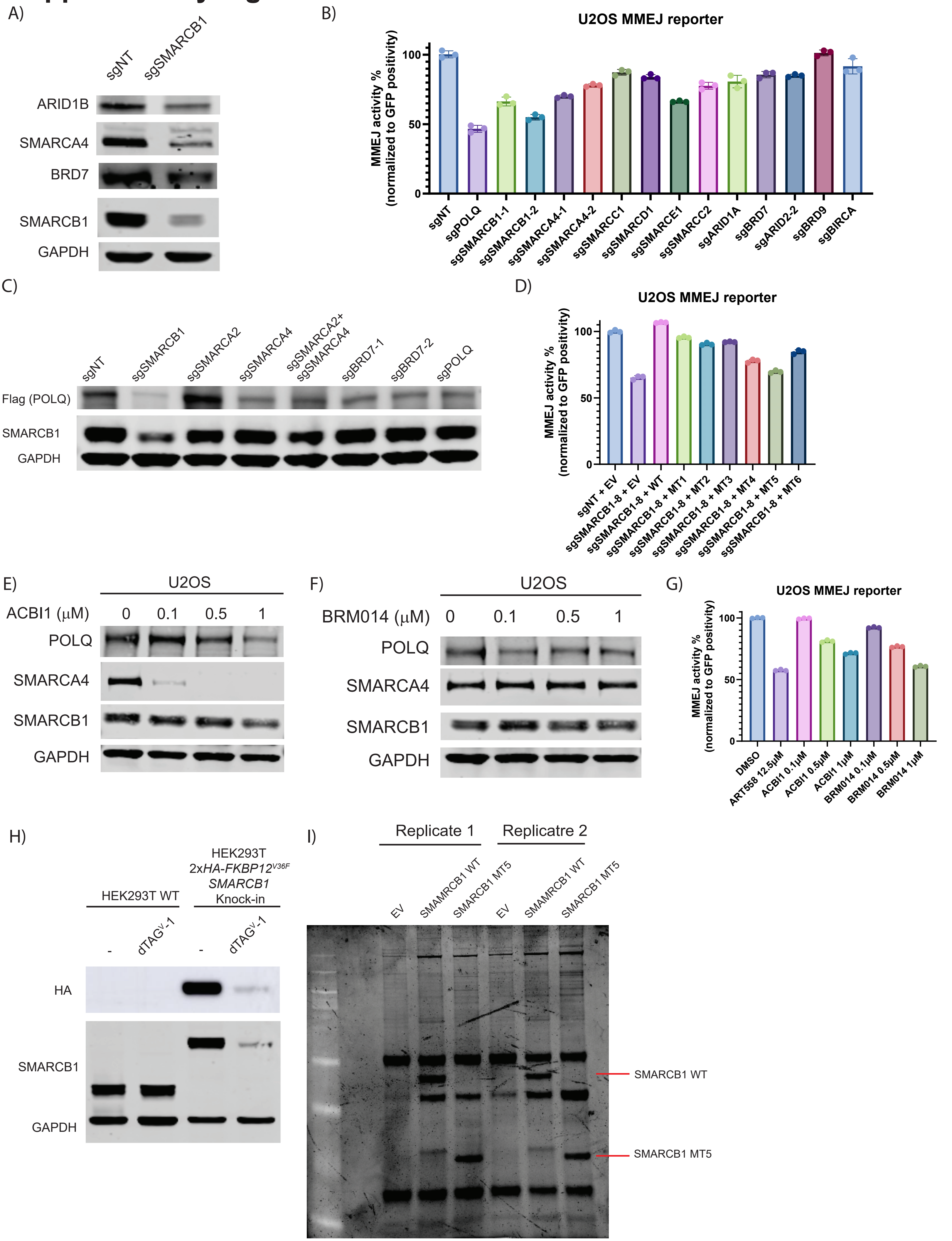

**Fig. S4. SMARCB1 regulates MMEJ activity and POLQ levels through cBAF/pBAF integrity.**

(A) Immunoblot of ARID1A, SMARCA4, BRD7, SMARCB1, and GAPDH in U2OS cells expressing sg*NT* or sg*SMARCB1*.

(B) MMEJ reporter assay in U2OS cells expressing the indicated sgRNAs targeting BAF complex subunits or *POLQ*.

(C) Immunoblot analysis of Flag-tagged POLQ and SMARCB1 in RPE *TP53$^{-/-}$ 3xFLAG-POLQ* knock-in cells expressing indicated sgRNAs. GAPDH serves as a loading control.

(D) MMEJ reporter assay in U2OS cells expressing an sgRNA targeting endogenous *SMARCB1* (but not the exogenous SMARCB1 cDNA), reconstituted with EV, WT SMARCB1, or truncation mutants (MT1–MT6).

(E-F) Immunoblot of POLQ, SMARCA4, and SMARCB1 in U2OS cells treated with (E) ACBI1 or (F) BRM014. GAPDH is shown as a loading control.

(G) MMEJ reporter assay in U2OS cells treated with ART558, ACBI1, or BRM014.

(H) Immunoblot showing inducible degradation of HA- FKBP12$^{F36V}$-SMARCB1 in HEK293T HA- FKBP12$^{F36V}$-SMARCB1 knock-in cells treated with dTAG$^{V}$-1. GAPDH was shown as a loading control.

(I) Oriole staining of V5 coimmunoprecipitates from HEK293T *HA-FKBP12$^{F36V}$-SMARCB1* knock-in cells after dTAG$^{v}$-1–induced degradation and transfection with EV, V5-tagged WT SMARCB1, or V5-tagged MT5 SMARCB1 (n = 2).

### Supplementary Figure 5

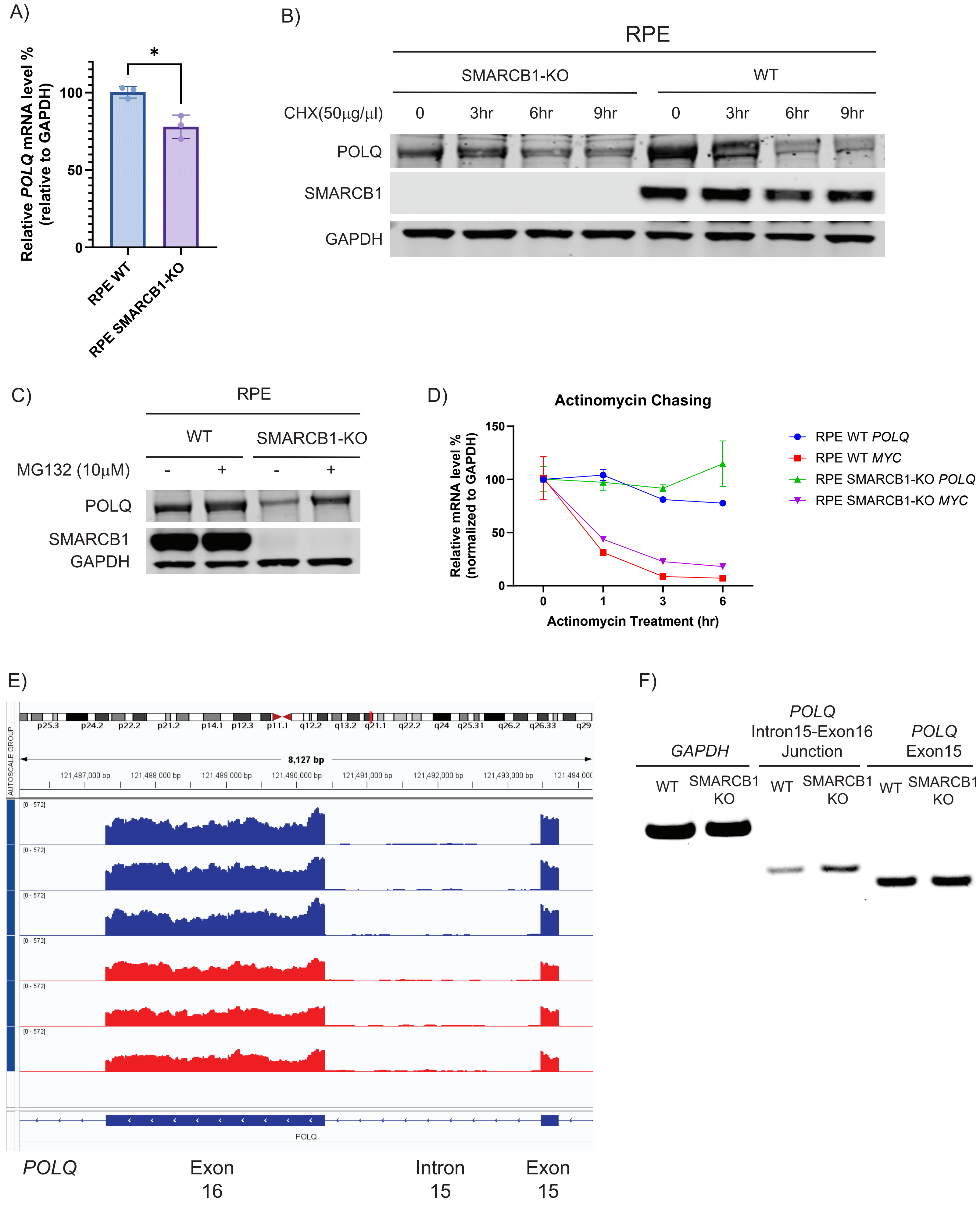

**Fig. S5. Impact of *SMARCB1* depletion on POLQ expression, protein stability, mRNA splicing, and mRNA stability.**

(A) *POLQ* mRNA expression in WT and *SMARCB1*-KO RPE cells measured by qRT-PCR, normalized to *GAPDH*. Data are shown as the mean ± SD. Statistical significance was calculated using Student's t test (*, $P < 0.05$).

(B) Cycloheximide chase assay of POLQ stability in WT and *SMARCB1*-KO RPE cells. Cells were treated with cycloheximide (50 µg/ml) for the indicated times.

(C) Immunoblot analysis of POLQ levels in RPE WT and *SMARCB1*-KO cells treated with or without the proteasome inhibitor MG132 (10 µM for 6 hr).

(D) Actinomycin D chase assay measuring *POLQ* and *MYC* mRNA stability over time in WT and *SMARCB1*-KO RPE $TP53^{-/-}$ cells.

(E) Genome browser view showing RNA-seq read coverage across *POLQ* intron 15 and its flanking exon 16 and exon 15 regions in wild-type (red) and SMARCB1-KO (blue) RPE $TP53^{-/-}$ cells.

(F) RT-PCR detection of the *POLQ* intron 15–exon 16 junction, exon 15, and exon 16 in RPE WT and *SMARCB1*-KO cells. *GAPDH* was amplified as a loading control.

### Supplementary Figure 6

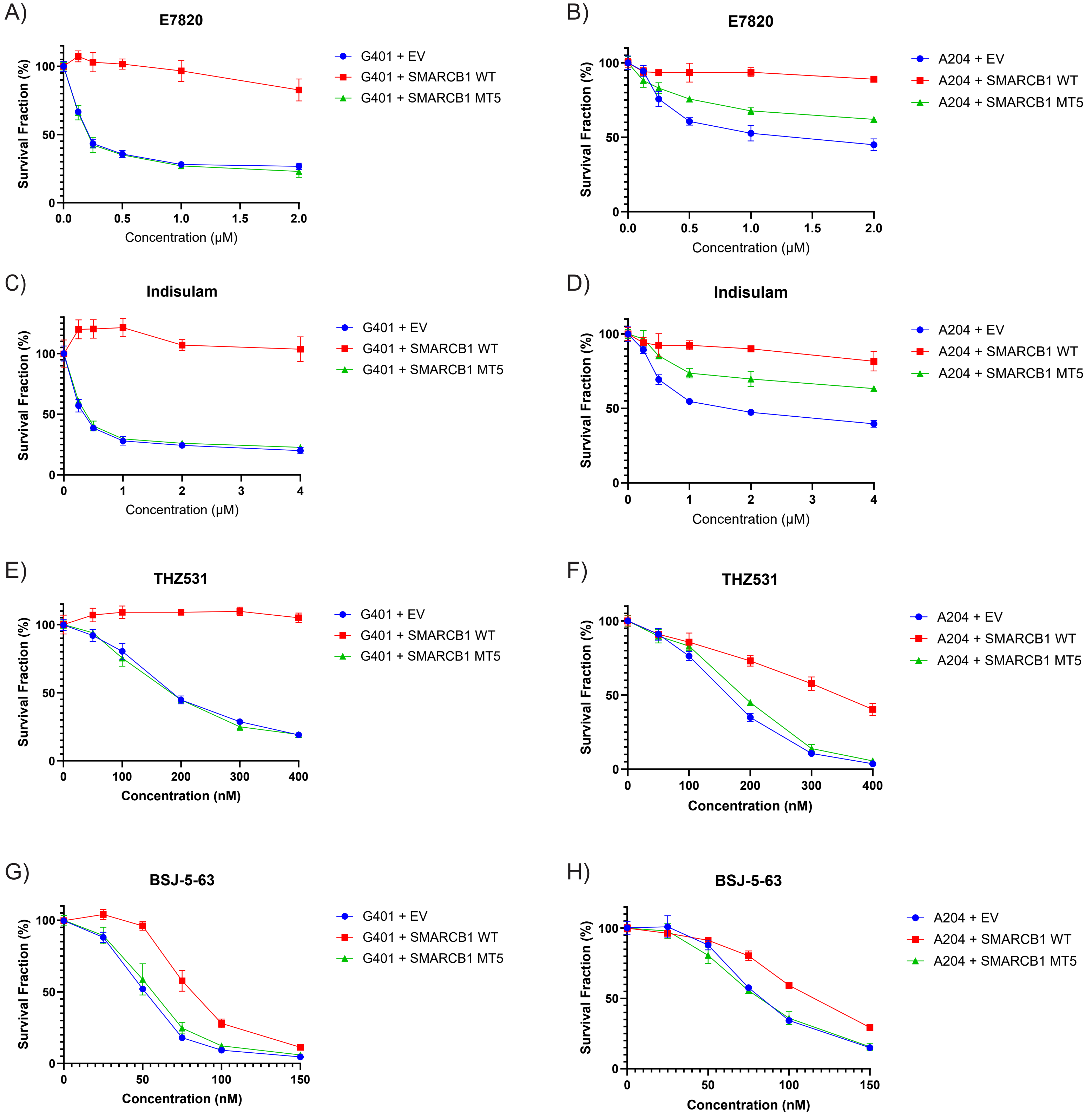

**Fig. S6. Rescue of SMARCB1 deficiency alters sensitivity to splicing inhibitors and RBM39 degraders.**

(A-B) Survival curves of (A) G401 cells and (B) A204 cells reconstituted with EV, WT SMARCB1, or MT5 and treated with indicated concentrations of E7820.

(C-D) Survival curves of (C) G401 cells and (D) A204 cells reconstituted with EV, WT SMARCB1, or MT5 and treated with indicated concentrations of Indisulam.

(E-F) Survival curves of (E) G401 cells and (F) A204 cells reconstituted with EV, WT SMARCB1, or MT5 and treated with indicated concentrations of THZ531.

(G-H) Survival curves of (G) G401 cells and (H) A204 cells reconstituted with EV, WT SMARCB1, or MT5 and treated with indicated concentrations of BSJ-5-63.
